# Single-molecule nucleosome spacing coordinates chromatin fiber interactions

**DOI:** 10.64898/2026.06.02.729033

**Authors:** Kaite Zhang, Maria Julia Maristany, Jan Huertas, Rosana Collepardo-Guevara, Vijay Ramani

## Abstract

Nucleosome spacing influences higher-order chromatin fiber organization *in vitro* but how this relates to cellular chromosome structure remains contentious. To address this, we developed Ligation Analysis of Single-molecule Sequence Interactions (LASSI), a single-molecule epigenomic method that combines proximity ligation with near-nucleotide resolution, long-read adenine methyltransferase footprinting. LASSI measures nucleosome spacing patterns on interacting segments of chromatin genome-wide in cells, quantifying coupled chromosome organization in 1D and 3D. Applying LASSI to mouse embryonic stem cells (mESCs), we discover a genome-wide structural pattern we term ‘fiber homotypy,’ where chromatin fibers with similar nucleosome spacing patterns interact more frequently in 3D. This pattern persists over long intrachromosomal distances in *cis* and interchromosomally in *trans*. Fiber homotypy negatively scales with genomic distance, though differently than contact probability, implying distinct mechanisms. It is further promoted by topologically associated domains (TADs), A/B compartments, and shared histone modification domains, suggesting instructive roles for each of these processes. Coarse-grain molecular dynamics simulations of chromatin fibers informed by LASSI reveal that the intrinsically heterogeneous spacing of nucleosomes along chromatin fibers in cells is a key regulator of homotypy. This variability tunes the free energy landscape of chromatin fiber interactions, promoting the compartmentalization of similar 1D chromatin structures via specific types of fiber-fiber interaction. Further linking compartmentalization and fiber homotypy, we demonstrate that loop extrusion antagonizes this phenomenon. Depletion of the cohesin subunit RAD21 in mESCs increases fiber homotypy genome-wide, while depletion of the cohesin unloader WAPL decreases fiber homotypy, consistent with effects seen on A/B compartmentalization. Our results demonstrate that fiber-fiber interactions driven by shared nucleosome spacing patterns instruct higher-order chromosome organization. Moreover, we show clear structural interdependence across cellular chromatin length-scales, likely tuned by processes ranging from nucleosome positioning to loop extrusion.

## INTRODUCTION

Cells in our bodies share largely identical genomes that vary in structure, gene expression, and function. This variability is regulated by the nucleoprotein complex chromatin. Chromatin is hierarchically and non-randomly organized across a range of length scales: from histone-containing nucleosomes, to ‘oligonucleosomes’ (*i.e.* nucleosomes successively spaced along DNA), to larger-scale structures (*e.g.* domains, compartments, and territories)(*1*, *2*). Though structure at each length scale impacts cell type specific nuclear functions(*3*, *4*), how structural patterns are coordinated across different length scales (for example, between 1D oligonucleosomes and 3D spatial domains) remains unclear.

Classical studies of chromatin suggested that the nucleosome spacing along DNA influences higher-order chromatin structure(*5*, *6*). Nucleosome spacing is computed as the average distance between nucleosomes along DNA (*i.e.* the nucleosome repeat length, or NRL). *In vitro* reconstitution has demonstrated stereotyped higher-order chromatin structural motifs, like folds with diameters of ∼30 nanometers (nm). These patterns arise from uniformly spaced NRLs at fixed values (∼147 + 10*n* bp, with *n* = 1, 2, 3, .. , ∼5), which promote ordered nucleosome packing within the fiber and give rise to ‘zig-zag’ chromatin structures(*6*, *7*). While these ordered structures are unlikely to stably persist in the nucleus(*8*, *9*), recent work has found that NRL also modulates the ability of chromatin to undergo liquid-liquid phase separation (LLPS) *in vitro*(*10*, *11*) by dictating local fiber structure and controlling the balance between intra- and inter-fiber interactions(*9*, *12*). Uniform NRLs of 147 + 10*n* sequester nucleosomes into strong intra-fiber stacking interactions, whereas deviations from this periodicity shift interactions toward inter-fiber contacts that promote phase separation(*9*, *12*). These *in vitro* phase separation studies showcase the fundamental biophysical principles that organize chromatin and other macromolecules in the nucleus(*13*, *14*), suggesting that NRL may influence higher-order chromatin organization *in vivo*.

Determining the link between NRL and higher-order chromatin organization in cells has remained challenging. *In vivo* studies have demonstrated substantial heterogeneity in chromatin fiber structure at the single-molecule level, at both levels of 1D nucleosome spacing and 3D chromosome organization. These studies (*e.g.* single-fiber nucleosome footprinting(*15–18*); single-cell Hi-C(*19*, *20*); single-nucleosome imaging(*21*, *22*); chromatin tracing(*20*, *23*); electron microscopy(*8*)) suggest extremely dynamic chromatin organization, even in transcriptionally inactive genomic regions. Accurately quantifying signal in this heterogeneous structural regime is further complicated by the known activities of dynamic motors on chromatin, like the SMC-family ATPase complex cohesin(*24*, *25*).

Here, we develop and apply an epigenome sequencing method – <u>L</u>igation <u>A</u>nalysis of <u>S</u>ingle-molecule <u>S</u>equence <u>I</u>nteractions (LASSI) – that addresses these limitations. LASSI measures single-molecule nucleosome spacing between physically proximal chromatin segments by combining single-molecule long-read footprinting with chromosome conformation capture (3C) technology(*26–28*). Resulting data quantify the extent to which nucleosome spacing and 3D genome organization converge in mammalian cells, with implications for NRL-mediated higher-order chromosome folding and its regulation by *trans* acting factors.

## RESULTS

### *G*enome-wide, simultaneous mapping of nucleosome spacing and chromatin contacts

We(*15*, *16*, *29–32*) and others(*17*, *33–37*) previously established high-resolution long-read methyltransferase chromatin footprinting to quantify nucleosome spacing at molecular resolution. LASSI (schematic shown in **Fig. 1A**) builds on this work, as well as previous attempts combining 3C with genomic footprinting(*38*, *39*), with key differences. LASSI is optimized for high (*i.e.* near-nucleotide) resolution footprinting of protein-DNA interactions on 2 or more interacting pieces of interacting DNA. As such, this protocol uses a six-cutter restriction enzyme (HindIII) to maximize the length of ligated chromatin segments. Moreover, crosslinking is carried out after methyltransferase footprinting, as even low amounts of crosslinker dramatically inhibit methylation efficiency. LASSI data is analyzed at the level of correlated signal on individual sequenced molecules and not with the purpose of generating 2D contact probability maps (as in *e.g.* Pore-C(*40*), CiFi(*41*), region-capture Micro-C [RCMC](*42*), or Micro-Capture-C [MCC](*43–45*)). Finally, unlike previous methods that call nucleosome spacing from high-resolution Micro-C data(*44*, *46*), which are limited by both the bulk aggregation of short-read sequencing and distance between nucleosomes (primarily capturing contacts between two neighboring nucleosomes), LASSI directly measures nucleosome positioning per molecule genome-wide. These conscious design decisions make LASSI uniquely suited for measuring kilobase-scale nucleosome spacing patterns on physically interacting chromatin fibers.

**Fig. 1:**
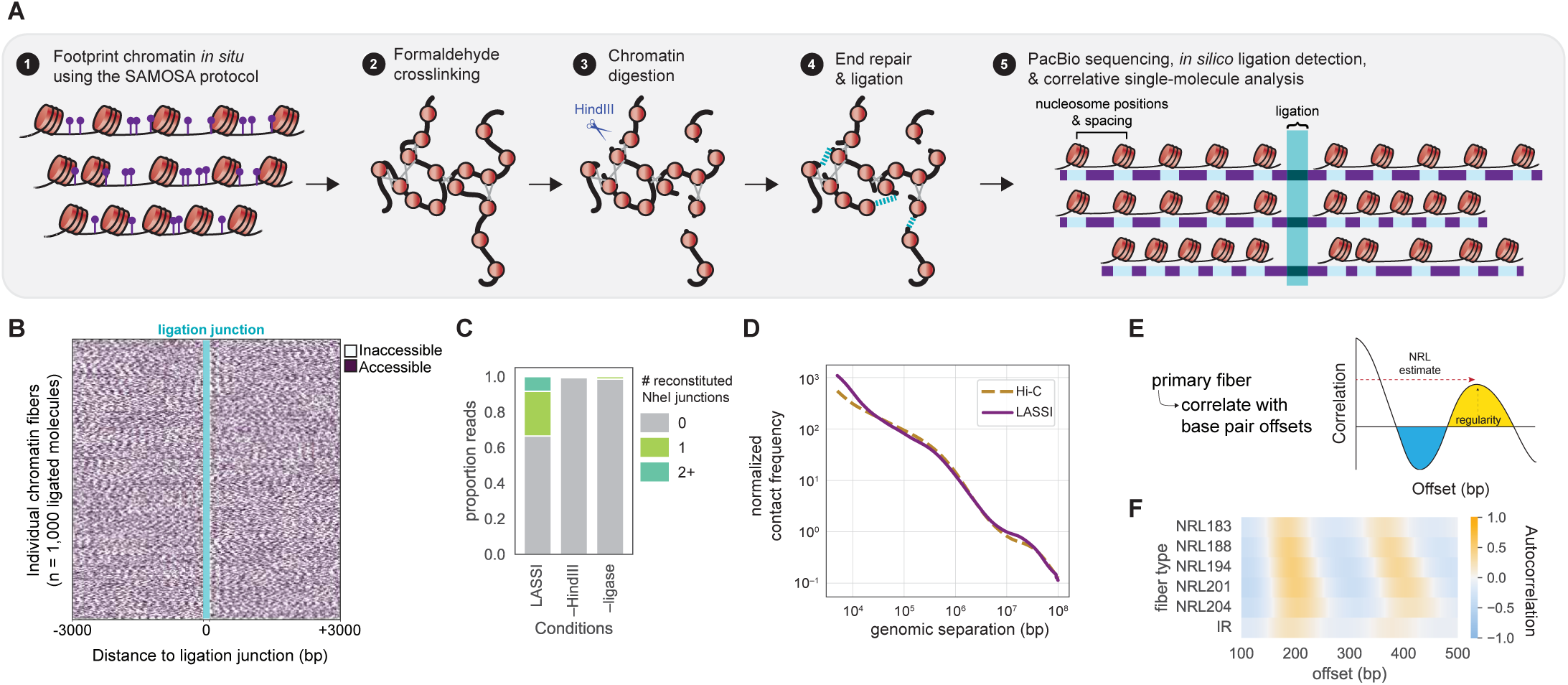
LASSI captures single-molecule nucleosome positioning on interacting chromatin fibers. **A.)** Overview schematic for the LASSI protocol. In nuclei, chromatin is first footprinted using the non-specific methyltransferase EcoGII. Then, chromatin is crosslinked, digested, repaired, and ligated in nuclei to form hybrid fibers that associate two disparate regions of the genome, spanning up to entire chromosomes. **B.)** Example of 1,000 random hybrid fibers, demonstrating footprinted chromatin out to 3 kb away from the ligation junction (teal). Regions that are inaccessible (white) are most commonly nucleosomes. **C.)** Proportion of reads with reconstituted ligation junctions, identified as NheI restriction sites, with either the complete LASSI protocol (which includes both HindIII digestion and T4 ligation), without HindIII, or without ligase. **D.)** Contact frequency decay by cis pair distance as measured by LASSI vs. comparable (6-cutter) Hi-C(*48*). **E.)** Schematic demonstrating autocorrelation analysis to determine fiber type. **F.)** Distribution of fiber types in wild-type mouse embryonic stem cells using LASSI.

We performed pilot LASSI experiments in mouse embryonic stem cells (mESCs). After performing the LASSI protocol, we sequenced ligated and footprinted DNA on the Pacific Biosciences sequencer and analyzed as previously(*16*, *29*) to call methyltransferase-inaccessible footprints. Sequencing reads fell into a mixed population of unligated, singly ligated, and multi-way ligation molecules, computationally identified by searching for reconstituted ligation junctions (accessibility of sampled molecules shown in **Fig. 1B**; **Fig. S1A-S1B**). Ligated molecules (hereafter ‘hybrid’ molecules; N = 976,460 interactions) were dependent on both restriction digestion and T4 DNA ligase (**Fig. 1C**; **Fig. S1A**), with observation of all four ligation orientations within data (**Fig. S1C**). Ligations were mainly intrachromomosomal (*i.e.* in *cis*), although interchromosomal (*trans*) interactions were observed with varying frequencies depending on nuclear extraction method (*trans* percentage 13.9% - 16.9% with digitonin (12% of reads for WT mESCs) and 29.9% - 42.1% with NP-40 across replicates; **Fig. S1D**). Finally, we observed contact probability decay curves in LASSI data comparable to multiple published Hi-C datasets(*4*, *47*, *48*) (**Fig. 1D, Fig. S1E**). These analyses validate LASSI as a *bona fide* proximity ligation assay.

We next confirmed the quality of LASSI nucleosome footprinting data with respect to the parent SAMOSA protocol. As previously, we performed single-molecule autocorrelation analysis and clustering (schematic in **Fig. 1E**) of individual chromatin segments. Doing so yielded 6 distinct fiber types that distill the heterogeneous chromatin footprinting signals into clusters sharing NRL and nucleosome regularity. Fiber type distributions observed from LASSI data were consistent with prior SAMOSA experiments in mESCs(*16*, *31*), with 5 clusters falling into regular patterns with approximate NRLs ranging from 183 to 204 base pairs (bp) and a single cluster capturing fibers with irregular (IR) nucleosome spacing (**Fig. 1F, Fig. S1F**). These experiments establish that LASSI simultaneously records nucleosome spacing and physical proximity of interacting chromatin fibers.

### Physically proximal chromatin fibers share nucleosome spacing patterns

We reasoned that if NRL-mediated chromatin fiber interactions instruct genome organization, LASSI should capture specific patterns of fiber type co-occurrence on ligated molecules. To test this hypothesis, we classified all hybrids by fiber type and calculated the odds that different fiber types would be captured together compared to chance (**Fig. 2A**). We observed a significant enrichment of self-self (*i.e.* ‘homotypic’) fiber type interactions, together with selective enrichment and depletion among specific self-nonself (*i.e.* ‘heterotypic’) fiber type pairs (**Fig. 2B, Fig. S2A-B**). Homotypic hybrids were universally significantly enriched (*e.g.* NRL204 - NRL204 Odds Ratio [O.R.] = 1.26; Storey’s *q* value = 6.18E-03). Significant enrichment of specific heterotypic hybrids was also observed, specifically between fiber types with similar NRLs (*e.g.* NRL183–NRL188; ι1NRL = 5; O.R. = 1.07; Storey’s *q* value = 2.98E-17). Conversely, heterotypic hybrids with larger differences in NRL were significantly depleted (*e.g.* NRL188–NRL204; ι1NRL = 16; O.R. = 0.838; *q* = 7.95E-03). These results demonstrate quantitative preferences in chromatin fiber interactions in mESCs, with interaction propensity increasing with similar NRL. We hereafter refer to this phenomenon as ‘fiber homotypy.’

**Fig. 2:**
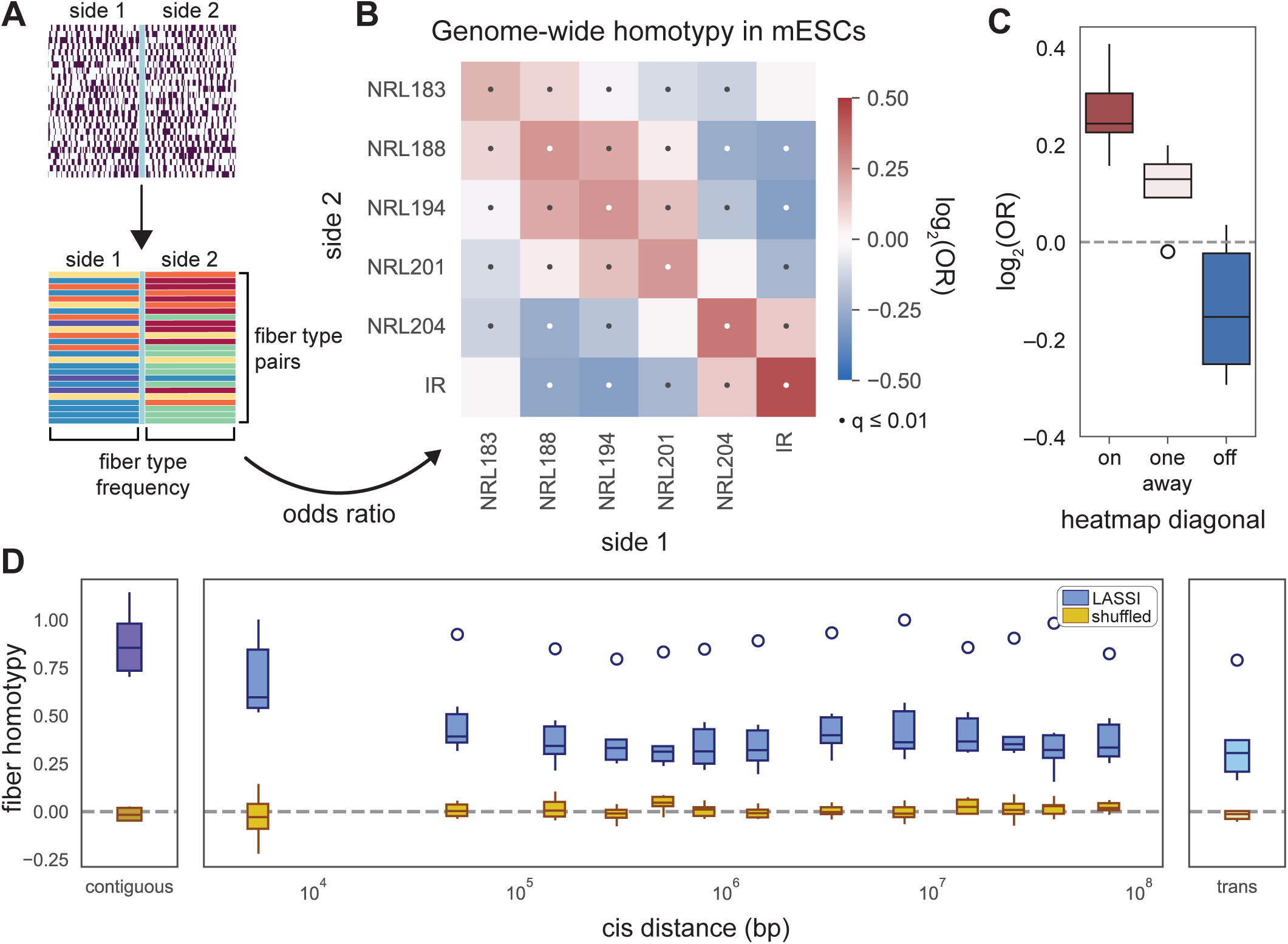
LASSI reveals genome-wide fiber homotypy in spatially proximal chromatin. **A.)** Schematic on how each side of hybrid fiber pairs (each row) are assigned a fiber type and frequency across the genome is calculated for each fiber type (each side of the junction), leading to odds ratio of fiber types found together in hybrids. **B.)** Odds ratio heatmap for wild type mouse embryonic stem cells (mESCs) of pairs of fiber types found in hybrid fibers, where the strong enrichment in self-associated fiber types (diagonal) is termed fiber homotypy. **C.)** Boxplot representation of heatmap in Fig. 2B, showing fiber homotypy as the “on” diagonal in red. **D.)** Fiber homotypy with relation to pair distance in *cis*, bookended by contiguous fibers (or non-digested, non-ligated fibers) and *trans* hybrid fibers. Fiber homotypy calculated from LASSI fibers is shown in blue; homotypy calculated from shuffled fibers per bin is shown in yellow.

We sought to understand how fiber homotypy varies as a function of genomic distance. To create a simple metric that captures fiber homotypy, we grouped all possible pairs in the interaction heatmap as identical (*i.e.* on-diagonal), similar (*i.e.* pairs one away from the diagonal), or dissimilar (*i.e.* off-diagonal). Odds ratios for each category were plotted together, offering a metric we termed fiber homotypy ‘score’ (**Fig. 2C**, red). We then assessed how scores varied as a function of *cis* interpair separation distance. To account for the nonuniformity of read depth across pair distances, we separated hybrid fibers into roughly equally sized bins (**Fig. S2C,D**) and included as further controls: i.) shuffled pairs from each bin, and ii.) scores computed for segments of unligated (‘contiguous’) reads. We also compared randomly sampled pairs of the same genomic distance away (**Fig. S2E**). We then visualized all scores (blue; **Fig. 2D**; alternate bin distribution in **Fig. S2F**) along with matched shuffled controls (yellow). As expected, shuffled controls demonstrated scores close to zero (**Fig. 2D**, **Fig. S2G-H**, right). Scores were highest for contiguous fibers, and the next highest scores observed for hybrid molecules with intrapair distances ≤ 10 kilobases (kb) (**Fig. 2D**, **Fig. S2G**, top left). Scores decayed to a baseline at ∼100 kb that was significantly elevated above random chance (*p* = 6.37E-258; Wald test). This baseline fiber homotypy score persisted up to the length of entire mouse chromosomes (∼10^8^ bp; **Fig. S2G**, bottom left) and for ligated molecules from different chromosomes (**Fig. 2D**). These data suggest that contact frequency influences (but does not completely explain) fiber homotypy.

Lastly, we tested whether fiber homotypy is limited to pairwise interactions or extends to higher-order chromatin interactions. The number of multi-way interactions (*i.e.* ‘C-Walks’)(*49*) in LASSI data decreases rapidly as a function of number of ligations (**Fig. S3A**); thus, we focused on C-Walks containing 3 segments exactly. To assess whether 3-segment C-Walks differ from 2-segment hybrids with respect to fiber homotypy, we computed odds ratios for the following conditions in three-way ligations: 1) all 3 segments have unique fiber types (maximum order of organization = 1; no fiber homotypy), 2) exactly 2 segments have the same fiber type (maximum order = 2; pairwise homotypy only), or 3) all 3 segments have the same fiber type (maximum order = 3; higher-order homotypy) (**Fig. S3B**). Fibers containing 3 ligated segments were even more likely to have 3 sides of homotypy than 2 sides alone (**Fig. S3C**), implying that fiber homotypy exists in neighborhoods of chromatin segments beyond interacting pairs.

### Increased fiber homotypy within TADs, compartments, and shared histone modification domains

We next assessed how fiber homotypy varies within 3D genome structural motifs. Genome structure can be categorized across length scales into units such as topologically associating domains (TADs) and A/B compartments. TADs are domains with increased contact frequency at kilobase resolution in *cis* (**Fig. 3A**), while compartments are typically defined by elevated contact frequency in *cis* and *trans* at megabase resolution (**Fig. 3B**). These structures are likely generated by opposing forces(*50*, *51*) and have distinct characteristics.

**Fig. 3:**
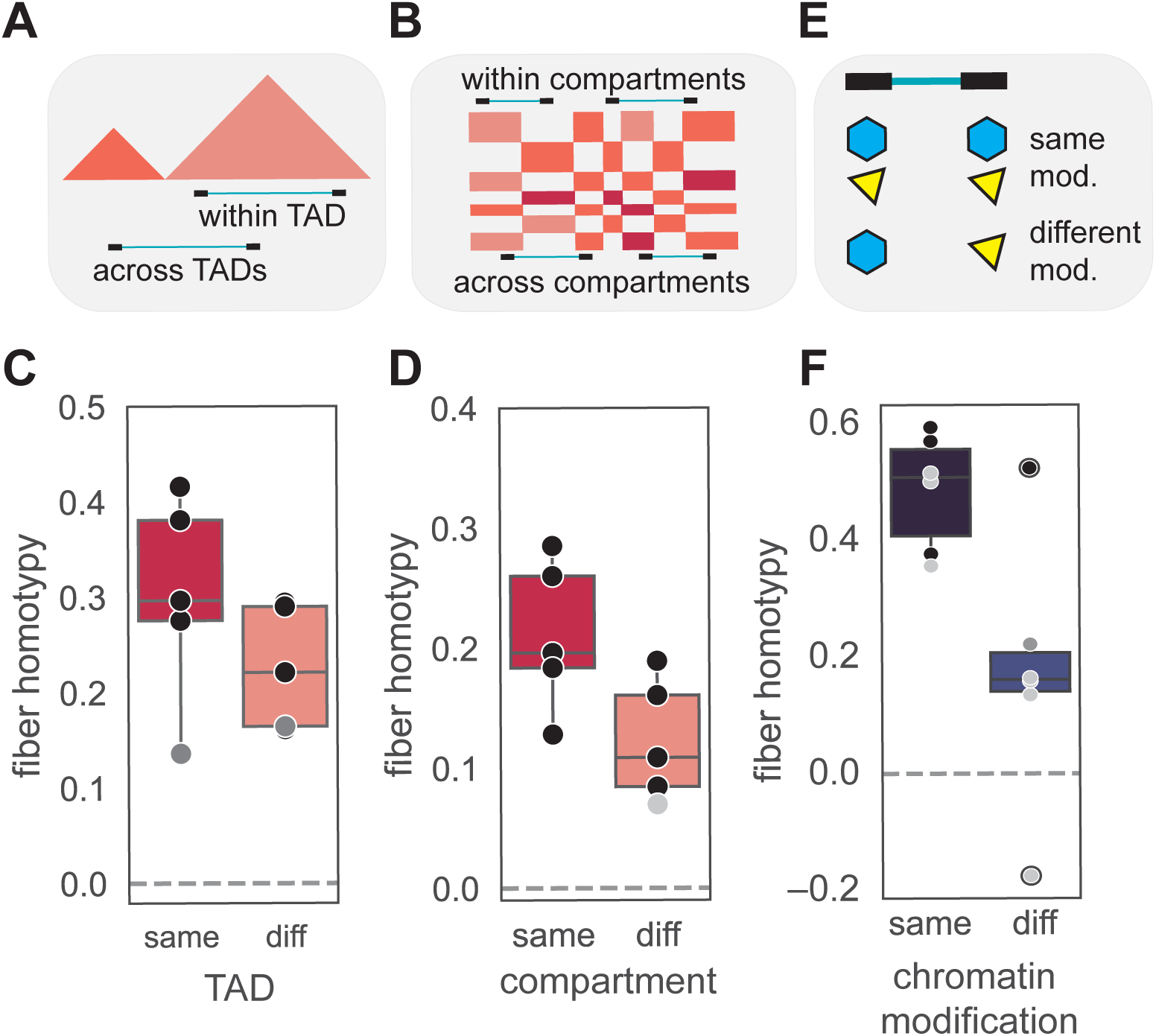
Fiber homotypy correlates with shared TADs, compartments, and histone modifications. **A.)** Schematic of how topologically associated domains (TADs) are typically depicted by Hi-C, where increased frequency of interaction at the kilobase scale is shown as intensity on a heatmap. Example ligation junctions (teal) between two sides of a hybrid fiber (black boxes) are shown, where the distance of the ligation is the same but either within the same TAD (“within”) or between different TADs (“across”). **B.)** Schematic of how megabase-scale compartments are typically depicted by Hi-C. **C.)** Fiber homotypy in distance-matched hybrid fibers, where homotypy is greater within the same TAD than between different TADs. Each dot is a fiber type. Black dots indicate q-value ≤ 0.01, dark grey dots indicate q-value ≤ 0.05, and light grey dots indicate q-value > 0.05. Irregular fiber types have been excluded. **D.)** Fiber homotypy in far cis (cis pair distance ≥ 5 Mb) and trans hybrid fibers, where homotypy is greater within the same compartment than between different compartments. Each dot is a fiber type. Black dots indicate q-value ≤ 0.01, dark grey dots indicate q-value ≤ 0.05, and light grey dots indicate q-value > 0.05. Irregular fiber types have been excluded. **E.)** Cartoon showing different histone marks, represented as polygons, where both sides of the hybrid fiber have the same or different histone modification. **F.)** Fiber homotypy in hybrid fibers that were at least 10 kb apart with either the same or different histone modification as defined by ENCODE ChIP-seq, where homotypy is greater with shared histone modifications. Each dot is a fiber type. Black dots indicate q-value ≤ 0.01, dark grey dots indicate q-value ≤ 0.05, and light grey dots indicate q-value > 0.05.

We first calculated fiber homotypy scores for hybrids within TADs(*4*). To mitigate the effect of genomic distance, we sampled molecules within and across TADs to match read depth across genomic distances (**Fig. S4**) and then performed comparisons. We observed significantly higher scores (*p* = 1.16E-02; Wald test) for hybrids within TADs compared to across TADs (**Fig. 3C**). We next performed a similar analysis at the level of A/B compartments. To reduce effects of genomic distance on both fiber homotypy and shared compartmentalization, we focused this analysis on hybrids spanning long distances in *cis* (≥ 5 megabases, Mb) and in *trans* (**Fig. S5A**). Homotypy scores were significantly higher within versus across compartments (**Fig. 3D**; *p* = 1.65E-04, Wald test). This correlation was robust to sampling across A versus B compartments (**Fig. S5B**), and randomly paired molecules drawn from the same compartments had a fiber homotypy score of 0 (**Fig. S5C**), suggesting the elevated fiber homotypy score within compartments is not simply an artifact of compartment-specific fiber structural patterns. We did, however, observe that cross-compartment homotypy was greater in *cis* compared to within-compartment homotypy in *trans* (**Fig. S5D**).

Compartmentalization is driven in part by the preferential self-association of similar chromatin states (*e.g.* histone modification / DNA modification domains)(*52*). Therefore, we next examined how fiber homotypy and structural patterns vary between and within chromatin fibers deriving from different chromatin states. We leveraged histone ChIP-seq data from the ENCODE project(*53*) to sort sequenced molecules by chromatin type (**Fig. 3E**). As with TADs and compartments, hybrid fibers with shared histone modifications demonstrated significantly higher fiber homotypy scores than those with different histone modifications (**Fig. 3F**; *p* = 8.46E-04, Wald test). This pattern was robust to domain-specific distribution differences in fiber type (**Fig. S6A-C**), genomic distance effects (**Fig. S6D**), and read depth (**Fig. S6E,F**). These analyses demonstrate that co-localization of chromatin within TADs, compartments, and shared histone modification domains promotes fiber homotypy beyond the genome-wide average.

Much of the genome is not clearly demarcated with histone modifications, and LASSI data captures many interactions between fibers with known modifications (‘modified’) and putatively ‘unmodified’ fibers. We next assessed whether spatial proximity between modified and unmodified chromatin fibers is sufficient to transmit state-specific nucleosome spacing information (**Fig. S7A**). We reasoned that if proximity to a chromatin state was sufficient to impart nucleosome spacing, then the unmodified ‘side’ of a modified-unmodified hybrid should bear modification-dependent spacing distributions. Intriguingly, despite the modified sides having state-specific nucleosome spacing patterns (**Fig. S6B**, **Fig. S7B** left), we observed no significant difference in fiber type distributions for the unmodified interacting pairs (**Fig. S7B** right). Next, we calculated fiber homotypy scores for hybrids for each mixed population, segregated by known histone modification. For most chromatin modifications, fiber homotypy scores approximated the genome-wide baseline (**Fig. S7C**). One key exception was H3K9me3-demarcated heterochromatin, which shows an off-axis pattern absent from other heterochromatic regions (**Fig. S7D**, top). This off-axis pattern persists after shuffling (**Fig. S7D**, bottom), suggesting it is an artifact of the fiber type distribution differences between H3K9me3 and putatively unmodified chromatin, possibly due to coverage biases for this annotated set of H3K9me3 regions in mESCs. Calculating normalized *z*-scores for actual versus shuffled hybrids restores the pattern of fiber homotypy (**Fig. S7E**). Taken together, these analyses suggest that spatial proximity to a histone modification domain does not alone explain the increased fiber homotypy scores observed within compartments or shared chromatin modification states.

### Fiber homotypy is thermodynamically driven by NRL heterogeneity and DNA content

We hypothesized there may be a biophysical basis for fiber homotypy, arising from intrinsic properties of nucleosomes along chromatin fibers. To test this hypothesis, we used molecular dynamics (MD) simulations. We applied our previously developed minimal coarse-grained chromatin model, which quantitatively captures the dependence of chromatin phase separation on NRL and reproduces NRL-dependent chromatin fiber structures within condensates seen by cryo-ET(*9*, *12*, *54*). Prior studies combining this approach with *in vitro* reconstitution demonstrated that uniformly spaced “147 + 10*n* bp” (‘canonical’) NRLs inhibit phase separation by locking nucleosomes in ordered intra-fiber interactions. Even small (±1 bp) deviations from canonical NRLs frustrate intra-fiber packing, shifting nucleosomes towards inter-fiber contacts and progressively promoting phase separation with optimal phase separation occurring at ±5 bp offsets(*9*, *12*). LASSI data (and our previously published single-molecule footprinting datasets(*16*, *29*, *31*, *32*)) demonstrate that chromatin fibers *in vivo* deviate considerably from canonical NRLs, with variability even beyond the limits of ±5 bp offsets around a single canonical spacing (**Fig. 4A**; **Fig. S8A**).

**Fig. 4:**
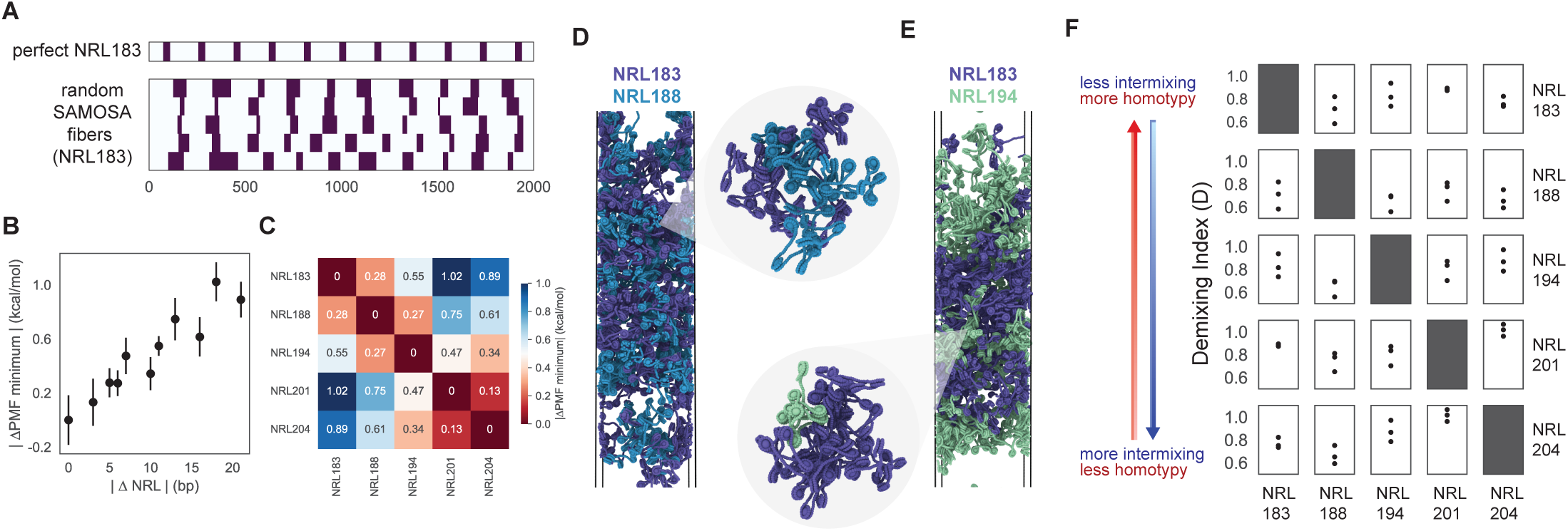
Molecular dynamics modeling recapitulates fiber homotypy found in vivo. **A.)** SAMOSA representation of idealized fibers with exactly 183 bp distance between nucleosome dyads compared to fibers of fiber type NRL183. **B.)** Binding free energies for idealized in vivo (IIV) fibers, plotted by the difference in fiber type-defined NRL. Error bars show ± 1 SEM. **C.)** Binding free energies for SAMOSA fibers as a heatmap for each heterotypic pair. Homotypic pairs have a |ΔPMF minimum| of 0. **D.)** Examples of fiber types NRL183 and NRL188, which tend to intermix (analogous to having less homotypy) in direct coexistence simulation. **E.)** Examples of fiber types NRL183 and NRL194, which tend to segregate in direct coexistence simulation (analogous to having more homotypy). **F.)** Demixing index for each pair in slab simulation following voxel analysis (see Methods and **Fig. S9**). Each dot is a replicate of a simulation of 60 SAMOSA-informed fibers.

To understand how NRL variability influences the physics of *in vivo* chromatin fiber structures, we performed simulations using fibers directly informed by LASSI data (*i.e. in vivo* or ‘IV’ fibers). We subsampled spacings from 200 13-nucleosome fibers for each fiber type observed (**Fig. S8B**) and simulated a single molecule’s behavior in space. These simulations demonstrate that IV fibers adopt highly irregular conformations sampling polymorphic structural ensembles that differ from those of control fibers with matching but regular NRLs (**Fig. S8C**). This confirms that deviation from regular NRLs is sufficient to expand the conformational phase space accessible to chromatin fibers.

We next explored how NRL influences the free energy landscape of fiber-fiber interactions *in silico*. The intrinsic heterogeneity of nucleosome spacing within each fiber type generates a large ensemble of possible 13-nucleosome fibers, each with distinct spacing patterns. Comprehensively calculating binding free energies for all IV fiber-fiber combinations across this ensemble of possible conformations is computationally intractable. Thus, we constructed idealized IV fibers (*i.e.* ‘IIV’ fibers) for each fiber type. IIV fibers use the fiber type NRL but remove DNA length-dependent torsional constraints, treating DNA as a flexible polymer. This allows us to recapitulate some of the structural heterogeneity of IV ensembles while systematically probing the impact of NRL on chromatin condensation.

We computed the free energy of all IIV fiber-fiber interactions across homotypic and heterotypic pairs (**Methods**). For idealized fibers, we previously observed that binding free energies exhibit an oscillatory dependence on linker length, reflecting two structural regimes: one in which linker lengths promote strong intra-fiber stacking but weak inter-fiber binding, and another in which they frustrate intra-fiber packing but inter-fiber interactions (lower binding free energies)(*12*). In stark contrast, binding free energies for IIV fibers lose oscillatory behavior, varying weakly and monotonically with NRL and reflecting their polymorphic structural ensembles. The total free DNA content of the fiber strongly influences interaction strength. Shorter NRL fibers exhibit stronger inter-fiber interactions, while longer NRL fibers exhibit weaker inter-fiber interactions, reflecting a growing imbalance between attractive nucleosomal contacts and repulsive free DNA. Heterotypic pairs thus incur a free-energy penalty, that increases with the difference in NRL (ι1NRL; **Fig. 4B**). Fiber pairs that colocalize experimentally correspond to those with the most similar interaction free energies in our simulations, *i.e.* the lowest penalty for heterotypic interactions (**Fig. 4C**). These simulations imply that fiber homotypy at least in part stems from thermodynamically favorable, nucleosome-mediated, fiber-fiber interactions constrained by per-fiber free DNA content.

Finally, we simulated how these thermodynamic principles manifest at the mesoscale. We performed direct coexistence simulations(*55*) of 50:50 heterotypic IV fiber mixtures. We initialized each system with 120 total fibers, each sampled from SAMOSA footprinting data, and characterized a metric analogous to fiber homotypy score, which we refer to as the demixing index(*56*) (**Fig. S9A**). In the demixing index, higher values indicate reduced compatibility between fiber types (*i.e.* disfavored heterotypic interactions and favored homotypic interactions). Excluding fiber type NRL204 (the second most heterogeneous fiber type after IR), direct coexistence simulations show that ΔNRL between fiber types is significantly correlated with selective association (**Fig. S9B**; Pearson’s *r* = 0.55, *p* = 0.016). Fiber pairs from more similar fiber types (*e.g.* NRL183–NRL188; **Fig. 4D**) mix readily, while those that are more dissimilar (*e.g.,* NRL183–NRL194; **Fig. 4E**) tend to demix (**Fig. 4F**). The demixing index for each heterotypic pair correlates with fiber heterotypy scores observed in *in vivo* LASSI data (**Fig. S9C**; Pearson’s *r* = 0.61, *p* = 0.008), with the exception of NRL204.

Taken together, our simulations demonstrate that NRL variability quantitatively regulates fiber-fiber interactions. Preferential associations emerge in regimes where different chromatin fibers compete for interactions, as small changes in NRL-dependent inter-fiber binding free energies are sufficient to selectively reorganize chromatin at the mesoscale.

### Loop extrusion antagonizes fiber homotypy

Loop extrusion by the cohesin holoenzyme spatially organizes chromatin through TAD formation(*57*, *58*) but also antagonizes A/B compartmentalization(*50*, *59*). Given the results of our minimal *in silico* MD simulations, we speculated that perturbing cohesin in cells might clarify links between fiber homotypy and A/B compartmentalization. If fiber homotypy and compartmentalization were linked, for example, then cohesin disruption would increase fiber homotypy genome-wide (**Fig. 5A**). Alternatively, cohesin depletion could lead to decreased fiber homotypy, implying other models, possibly involving chromatin constraint by cohesin(*60*). To distinguish between these models, we performed LASSI experiments using auxin-inducible degron system to deplete the cohesin subunit RAD21 in mESCs. We selected two time points (90 and 210 min.) to understand dose-dependent effects (**Fig. 5B**; **Fig. S10A**), and then measured fiber homotypy using LASSI (digitonin permeabilization). Depletion of RAD21 led to a significant increase in homotypy score (**Fig. 5C**); this increase was most pronounced for *cis* pairs between 100 kb and 1 Mb (**Fig. S10B**), the distances at which contact frequency changes most by P(*s*) analysis of LASSI data (**Fig. S10C**). These results suggest that cohesin negatively regulates fiber homotypy.

**Fig. 5:**
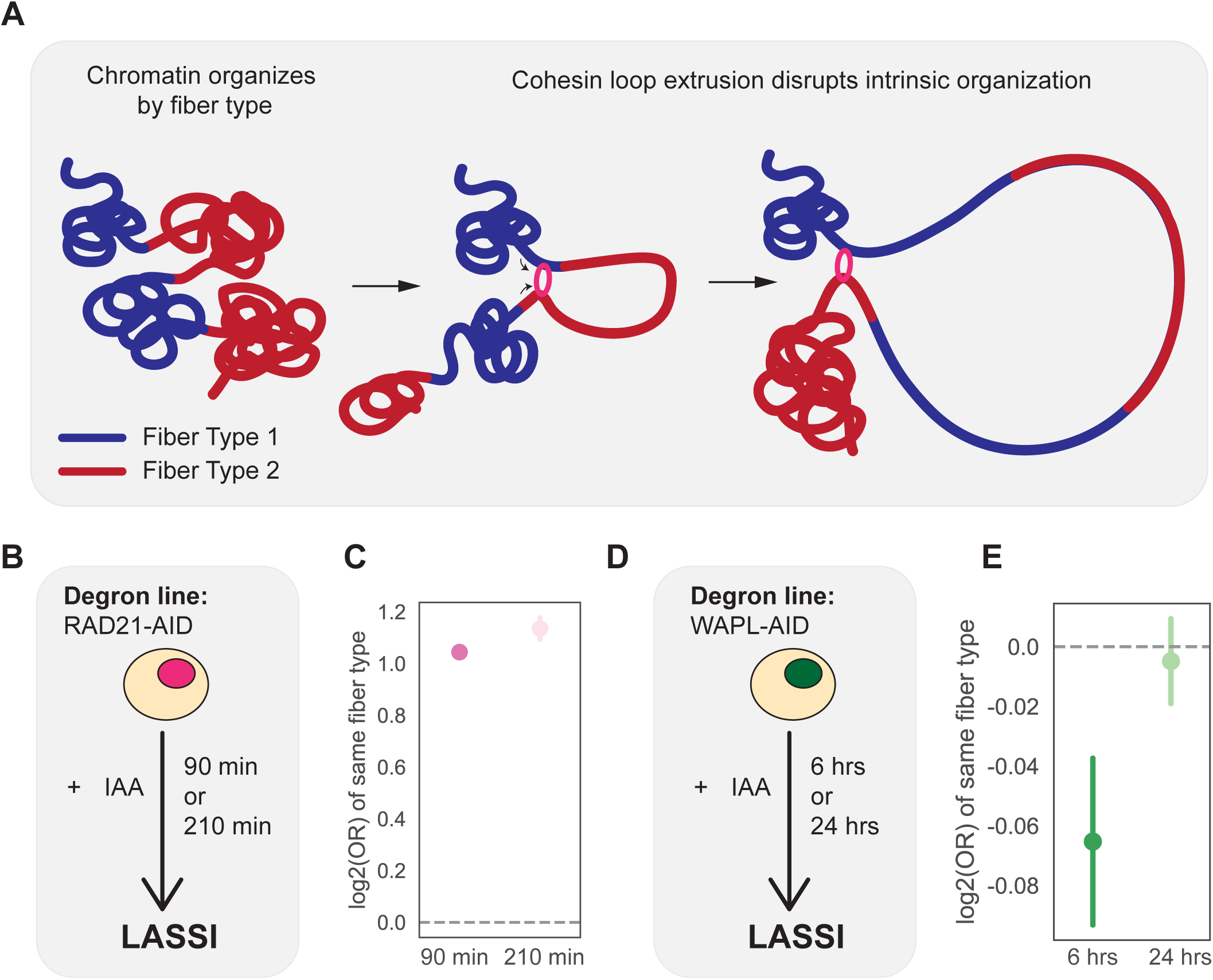
Ablating loop extrusion increases fiber homotypy while overextrusion decreases homotypy. **A.)** Schematic showing how otherwise self-organized fiber types (red and blue) may be disrupted by cohesin (pink loop). **B.)** Schematic for depletion of RAD21. **C.)** Log_2_ odds ratio of identical fiber types found in hybrid fibers under RAD21 depletion (either 90 min or 210 minutes) when compared to DMSO condition. p-value < 0.01. **D.)** Schematic for depletion of WAPL. **E.)** Log_2_ odds ratio of identical fiber types found in hybrid fibers under WAPL depletion (either 6 or 24 hours) when compared to DMSO condition. p-value = 0.08.

A model in which cohesin disrupts self-similar fiber interactions further predicts that hyperextrusion will decrease fiber homotypy scores genome-wide. To test this hypothesis, we performed a second depletion experiment targeting the cohesin unloading factor WAPL (**Fig. 5D**). We depleted WAPL in mESCS in biological duplicate for 6 hours and 24 hours and carried out LASSI as above. WAPL depletion led to the expected increase in contacts at distances > 1 Mb in the corresponding P(*s*) curve, indicating that depletion was successful (**Fig. S10D,E**), and led to a significant, albeit modest, decrease in homotypy score (**Fig. 5E**). Fiber homotypy score returned to baseline after 24 hours, suggesting steady-state equilibration of fiber type dependent interactions.

## DISCUSSION

### Single-molecule nucleosome spacing patterns are spatially coordinated in cells

We describe LASSI, a method that measures how single-molecule chromatin structure varies on physically proximal chromatin segments in cells. 3C and related methods measure contact frequency, at either increasingly granular resolutions(*25*, *44*, *61*) or increasing levels of higher-order conformation(*40*). LASSI instead measures the relationship between 1D nucleosome spacing and 3D spatial organization of kilobase-scale molecules, which has previously only been possible *in vitro*(*9*, *10*, *12*, *62*). LASSI is not without limitations, however. The approach can only capture physically proximal chromatin fibers and is blind to the background of distal chromatin fibers within the same nucleus (unlike single-cell, single-molecule approaches(*63*, *64*)).

Leveraging the combined power of proximity ligation and methyltransferase footprinting in cells, we demonstrate that spatially proximal chromatin is significantly more likely to share nucleosome spacing patterns, a phenomenon we term ‘fiber homotypy’ (**Fig. 6A**). Fiber homotypy is maximal within ∼100 kb in genomic distance but importantly remains present at low levels in far *cis* and in *trans*. Fiber homotypy is higher within TADs and compartments, suggesting that the spatial constraint of chromatin within these structural units coordinates 1D chromatin structure. We propose that this phenomenon reflects a ‘super-structure’ underpinning genome organization, that coordinates chromatin structure across length scales (**Fig. 6B**). We stress, however, that the direction of influence remains unclear. Is proper 3D spatial segregation of chromatin in nuclei solely instructive of 1D nucleosome spacing? Or is 3D spatial segregation a byproduct of the 1D chromatin fiber structure? Future studies using complex *in vitro* reconstitutions combined with genomics (*e.g.* single-molecule footprinting(*15*, *30*, *65*) or Hi-C(*62*) will be especially instructive in this regard.

**Fig. 6:**
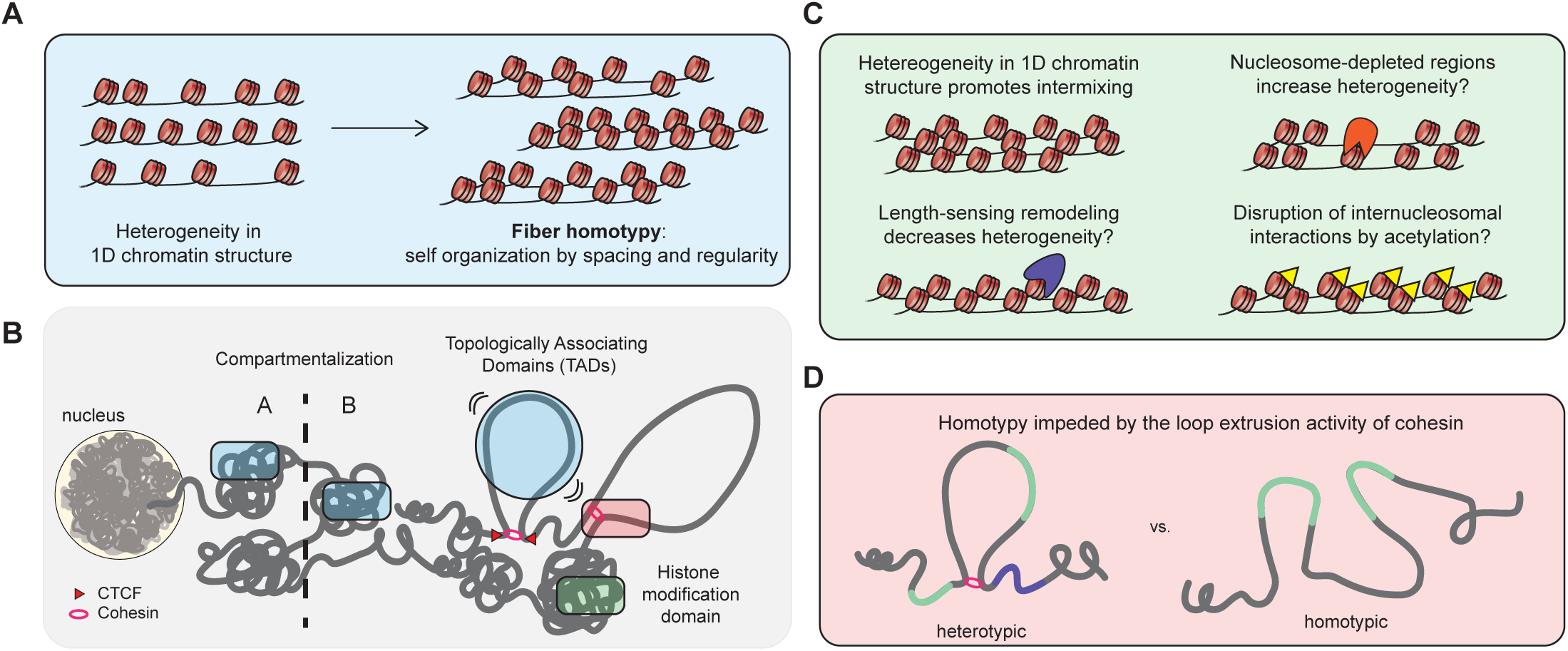
Chromatin spatial organization is multifaceted and involves linear chromatin structure. **A.)** Despite heterogeneity in nucleosome spacing (or 1D chromatin structure), chromatin in the nucleus associates with other chromatin that are similar in nucleosome spacing and regularity. **B.)** Classic overview of 3D genome organization at a range of scales. Translucent blue, green, and red overlays indicate how fiber homotypy may relate to these other phenomena, corresponding to (A), (C), and (D) respectively. **C.)** Illustrations of how heterogeneity is crucial to intermixing as revealed by MD simulations (top left) as well as speculation on how 1D nucleosome spacing may be tuned in nuclei to affect 3D organization. **D.)** Cartoon for the disruption of genome-wide fiber homotypy by the loop extrusion activity of cohesin.

### Single molecule chromatin structure regulates chromatin intermixing *in silico*

MD simulations reveal that distinct patterns of chromatin intermixing arise from NRL heterogeneity. Within this regime, interaction free energies are tuned by the total free DNA content of the fibers and govern mixing behavior, such that fibers with similar NRLs interact more strongly. *In vitro* studies have shown that nucleosome–nucleosome interactions, particularly intermolecular contacts between nucleosomal faces, are central to chromatin phase separation(*9*, *12*). The orientation of nucleosomal faces is entirely dependent on the NRL, and *in vivo* heterogeneity in NRL directly disrupts these geometric constraints, resulting in structurally irregular chromatin fibers that sample a broad conformational space. Future studies combining the experimentally-constrained MD framework established here with systematic perturbation of *e.g.* ATP-dependent chromatin remodeling enzymes will establish how this broad conformational space drives cell-type specific patterns of genome organization.

Importantly, we stress that as with any *in silico* experiment, the lack of nuclear context must also be considered. Our data suggest that 1D spacing can drive 3D conformational patterns, but these observations do not incorporate post-translationally modified histones, transcription factors, and other critical features observed in cells (*e.g.* HP1(*66*), speckles(*67*), transcription hubs(*14*)). Fully determining how nucleosome spacing, nucleosome biophysics, and the nuclear *trans* environment regulate chromatin conformation remains an important future direction (**Fig. 6C**) that will require further experimentation and modeling.

### Nucleosome spacing patterns contribute to chromosome compartmentalization

Integrating multiple lines of evidence from our study and others(*9*, *12*, *21*, *30*, *62*), we propose that fiber homotypy is a quantitative contributor to the long-observed pattern of chromosome compartmentalization. Our LASSI data and computational simulations together confirm that nucleosome-nucleosome interactions promote selective association of fibers on the basis of single-molecule fiber structure, adding to the growing list of mechanisms by which associative chromatin interactions may occur in nuclei (*e.g.* oligomerization of heterochromatin proteins(*66*, *68*), dimerization of LIM domains(*69*), condensation of chromatin-associated proteins via low-complexity protein sequences(*70*)). Our chemical genetic LASSI experiments further support this model by demonstrating an antagonistic relationship between loop extrusion and fiber homotypy (**Fig. 5E**, **Fig. 6D**), in alignment with results showing similar antagonism between loop extrusion and compartmentalization(*50*, *59*).

Future work clarifying the link between fiber homotypy and compartmentalization must focus on disentangling the direct effects of nucleosome spacing on chromosome folding versus other regulatory effects. Nucleosome remodeling complexes act redundantly(*71*) and can influence chromosome structure and cellular function in multiple ways(*3*, *72*). Testing this model in the future will require the systematic and combinatorial depletion of mammalian chromatin remodeling enzymes, followed by LASSI experiments.

### “Gelation by nucleosome” as an organizing principle for genomes

Virtually all biological systems must preserve liquid-like environments to carry out essential biochemical reactions. Failure to do so can lead to pathological aggregation of both nucleic acid(*73*) and protein(*74*). The results of our study, integrated with recent work from both us and others(*2*, *50*, *59*, *75*), lead us to propose a model of “gelation by nucleosome.” Chromatin fibers assembled *in vitro* on mammalian sequences condense but demonstrate minimal internal dynamics(*30*), especially when compared with condensates formed from chromatin assembled on synthetic sequences(*10*), but chromatin in dividing cells is far more dynamic. We speculate that nucleosome spacings present on native DNA sequences intrinsically lead to less dynamic, more crosslinked chromatin, which are then resolved by active processes to enable essential nuclear functions. By controlling the relative ‘gelation’ of particular chromatin states, motor activity itself may coordinate genome function across length-scales and drive cell-type-specific regulatory programs(*76–78*).

## Acknowledgements

We thank members of the V.R. laboratory, Elphège Nora, and Erika C. Anderson for discussion, experimental suggestions, and comments on the manuscript and T. Tolpa for generating schematics used in the figures. K.Z. was partially supported by the Hillblom/Bakar Aging Research Institute. This work was partially supported by NIH grants U01-DK127421 and DP2-HG012442 to V.R. V.R. acknowledges generous support from the Searle Scholars Award and the W.M. Keck Foundation.

## MATERIALS AND METHODS

### Cell culture

Mouse embryonic stem cells (mESCs) were grown on tissue culture plates (CellTreat 229106) after coating with 0.2% gelatin. Standard media contained DMEM + Glutamax (ThermoFisher 10566-016), 15% Fetal Bovine Serum (Cytiva SH30071.03), 1X non-essential amino-acids (ThermoFisher 11140-50), 1 mM sodium pyruvate (ThermoFisher 11360-070), 59 µM 2-Mercaptoethanol (BioRad 1610710XTU), 1 µM PD0325901, 3 µM CHIR99021, and 1X Leukemia Inhibitory Factor (purified and gifted by Barbara Panning Lab at UCSF). mESCs were fed with fresh media every day and passaged every 2 days using TrypLE Express Enzyme (1X) (ThermoFisher 12605010). Degron lines (RAD21-AID line EN272.2 and WAPL-AID line CMW184) were generously gifted from the Elphège Nora lab and Elzo de Wit lab respectively.

### Nuclei extraction and cell permeabilization

Nuclei were extracted using NE1 buffer (20 mM HEPES pH 7.5, 10 mM KCl, 1 mM MgCl_2_, 0.1% Triton X-100, 20% glycerol, 1X protease inhibitor) by incubation on ice for 10 minutes and washed in 1X Buffer M (15 mM Tris-HCl pH 8, 15 mM NaCl, 60 mM KCl, 0.5 mM spermidine) before proceeding with footprinting. Alternatively, cells were permeabilized in 1X Buffer M with 0.05% digitonin for 30 minutes on ice, before resuspending in 1X Buffer M + 0.05% digitonin for footprinting.

### Ligation analysis on single-molecule sequence interactions

#### Experimental protocol

Chromatin was footprinted using the SAMOSA protocol as previously described(*15*, *16*, *31*, *32*). Buffer M was washed out with 1X PBS and cells were fixed in 1% PFA (made fresh from Thermo Scientific 28906) in 1X PBS for 10 minutes with rotation. Fixation was quenched by spiking in fresh 1 M glycine (Sigma-Aldrich G7126) to a final concentration of 125 mM, incubating for 5 minutes at room temperature and then on ice for 15 minutes. Cells were washed in 1X NEBuffer r2.1 and then resuspend in 0.9 volume 1X NEBuffer r2.1 with 0.1 volume 1% SDS, incubated for 10 minutes at 65°C, and quenched with 0.11 volume 10% Triton X-100 for 10 minutes at 37°C. Final resuspension was in 1X NEbuffer r2.1 for overnight digestion by 80 U HindIII per million input cells. Chromatin was end repaired by adding end repair mix (0.15 mM dNTPs, 0.1 U/µL Klenow, 1 mM DTT) and incubating at 37°C for 45 minutes. Ligation proceeded by adding ligation mix (1X T4 DNA ligase buffer with ATP, 0.83% Triton X-100, 0.1 mg/mL BSA, and 50 Weiss U T4 DNA ligase per million cells) and incubating at room temperature for 3 hours. For DNA extraction, 10 µL RNAse A (Thermo Scientific EN0531) was added and chromatin incubated for 10 minutes at 37°C, followed by adding proteinase K mix (to a final concentration of 0.93 mg/mL proteinase K and 1.12% SDS) and incubation overnight at 65°C. Equal volume phenol:chloroform:isoamyl alcohol (Sigma-Aldrich 77617) was added and vortexed briefly to mix. After centrifugation at max speed (≥ 13,000 g) for 2 minutes at room temperature, the upper aqueous phase was transferred to a new tube, to which 0.1 volume 3M sodium acetate, 1 µL GlycoBlue (Thermo Scientific AM9516), and 3 volumes ice cold 100% ethanol was added. Extraction was incubated overnight at -20°C or for 2 hours at - 80°C, then centrifuged for 30 minutes at 4°C and max speed (≥ 13,000 g). Remaining pellet was washed with 80% ethanol, centrifuged for 2 minutes at the same temperature and speed, and then supernatant removed and pellet allowed to dry for 10 minutes. DNA was resuspended in 50 µL PacBio elution buffer. PacBio libraries were prepared using SMRTBell Prep Kit 3.0 and sequenced on the Sequel II or Revio sequencer using Polymerase Binding Kit 3.2.

#### Computational pipeline

Successful ligations (hereafter “hybrid fibers”) were identified by their reconstituted ligation junction AAGCTAGCTT. Hybrid fibers were identified pre-alignment, split by the center of the ligation junction, and then each “side” was aligned separately. Footprinting was called as previously described(*15*, *16*). Because footprinting was determined on pre-split, pre-alignment sequences, custom scripts were used to properly split and match footprints to sequences. Additionally, to account for any artifacts that might occur at the ligation junction, the 75 base pairs closest to the junction were omitted for downstream processing, including fiber type analyses using autocorrelation and Leiden clustering. For downstream analyses, read IDs (which include a sequencing ID and zero-mode waveguide (ZMW) number) with a zero-indexed split number (e.g., 0 for the first segment, 1 for the second segment, and so on) were used to identify unique ligations, and read information (including alignment) were stored in the HDF5 format.

Each unique ligation event was considered for downstream analyses. Odds ratios were calculated using the Fisher’s exact test. Z-scores were calculated using the formula 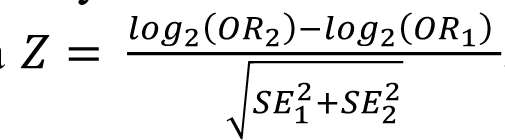 Significant differences between collections of odds ratios were calculated using a Wald test in R.

For calculating fiber homotypy score for Fig. S5, far cis consists of fibers that were ≥ 1Mb apart.

#### Hi-C processing

Publicly available Hi-C data(*47*, *48*, *60*, *79*) were retrieved from GEO, aligned to mm10 or hg38 as appropriate, and processed for plotting P(s) curves using pairtools(*80*), cooler(*81*), and cooltools(*82*). A custom script was used to convert LASSI data to the .pairs format and P(s) plotted using cooler and cooltools. Due to the differences in sequencing depth between Hi-C experiments, as well as between Hi-C and LASSI experiments, contact frequency was normalized by the mean contact frequency per sample for comparably plotting P(s) curves for all experiments.

Topologically associating domains and A/B compartments in wild type mESCs were retrieved from Ref(*4*).

#### Chromatin modification processing

All histone modification domains were retrieved from ENCODE(*53*) except for H2AK119Ub1, which was retrieved from Ref(*83*). Domains sourced from ENCODE were processed as previously(*15*). Histone modification peaks were overlapped with each molecule (or side, after LASSI processing) and molecules containing more than one type of modification were excluded.

### Western Blots

Cells were lysed in RIPA lysis buffer (150 mM NaCl, 0.1% Triton X-100, 0.5% sodium deoxycholate, 0.1% sodium dodecyl sulfate, 50 mM Tris-HCl pH 7.6, 1X protease inhibitor (Roche 04693159001), 0.1 mM PMSF (Sigma-Aldrich 10837091001) added fresh) at a concentration of 100 µL per million cells for 30 minutes on ice. Lysate was spun at max speed (≥ 16,000 g) for 10 minutes at 4°C and the clarified lysate (supernatant) was taken for blotting. Protein concentration was measured by BCA (Thermo Scientific 23227) and 20 µg of lysate was used per lane. Lysate was mixed with NuPAGE LDS Sample Buffer buffer (Thermo Scientific NP0007) on an 4-12% Bis-Tris gel (Thermo Scientific NP0321) in 1X MOPS SDS Running Buffer (Fisher Scientific NP000102) at 100 V for 10-15 minutes and then at 150 V for 60-90 minutes. Protein was then transferred to 0.45 µm PVDF (first rinsed in 100% methanol and then pre-soaked in transfer buffer) using 1X Novex Transfer buffer (Fisher Scientific NP00061) overnight in cold room at 30V. Successful transfer was confirmed with Ponceau Red, after which membrane was blocked using 5% milk in 1X PBST (0.1% Tween-20 in 1X PBS) for 30 minutes at room temperature. Primary antibody was diluted according to manufacturer’s instructions and incubated overnight at 4°C, after which membrane was washed three times in 1X PBST prior to incubation with secondary antibody diluted according to manufacturer’s instructions for 1 hour at room temperature. Membrane was washed three times in 1X PBST prior to imaging. Antibodies used: anti-WAPL (Proteintech # 16370-1-AP), anti-RAD21 (Abcam # ab154769), anti-vinculin (Sigma-Aldrich V9264), and anti-H3 (Cell Signaling Technology # 4499).

### Molecular dynamics simulations

#### Molecule sampling

LASSI hybrid fibers from 1 replicate were filtered for read length (≥ 2.5 kb), number of linker regions (≥ 12), and footprint size distributions (90-250 bp for all footprints on array). From this set, 200 fibers from each fiber type were randomly sampled as a pool of molecules for modeling, from which each replicate simulation used 60 molecules. For modeling, 12 consecutive linker lengths were sampled per each fiber and modeled with ideal nucleosomes, with an expected footprint size of 147 bp.

#### PMF Simulations

Potential of mean force (PMF) profiles were calculated using umbrella sampling as implemented in the COLVARS module(*84*). To improve configurational sampling within each umbrella window, we combined this approach with temperature-replica-exchange molecular dynamics (T-REMD). Replica exchange was performed independently within each umbrella window; no exchanges were allowed between different windows. For each window, 16 replicas were simulated over a temperature range of 300–600 K. The standard Metropolis criterion was applied for replica exchanges, with the biasing potential included in the total energy.

To mimic structural disorder, we removed DNA torsional constraints. For inter-fiber chromatin interactions, the collective variable was restrained using a harmonic bias with a force constant of 0.002 kcal mol⁻¹ Å⁻¹. A total of 16 umbrella windows were distributed evenly across the range 0–600 Å.

PMFs were reconstructed from the umbrella sampling data using the weighted histogram analysis method (WHAM)(*85*). All simulations were repeated independently 5 times. Reported PMFs values correspond to the average over these replicates.

#### Direct Coexistence simulations

To compute the phase diagram of 13-nucleosome chromatin, we employ the direct coexistence method(*55*, *86*) using 125 independent 13-nucleosome chromatin arrays at fixed salt concentration(*12*, *54*). The initial structures of the chromatin array are generated from T-REMD simulations at the corresponding salt concentration.

In a direct coexistence simulation, both the dilute and condensed liquid phases are placed within a single elongated simulation box. The simulation proceeds until both phases reach equilibrium at their respective coexistence densities. Once equilibrium is established, the coexistence densities can be determined by averaging the density profile along the long axis of the simulation box, with the center of mass held fixed.

Once equilibrated, the simulations were run for approximately X million timesteps, and coordinate snapshots were recorded every X timesteps. This is greater than the correlation time of our model, which is equal to 4000 timesteps (1 timestep = 100 fs). The simulation box dimensions were 1200 Å x 1200 Å x 12000 Å.

#### Correlation analysis

To compute the local correlation, or demixing, of the mixed simulations where two chromatin fiber species were presented, we adapted the method described in Ref(*56*).

The simulation box was divided into equal-sized voxels, and coordinates were wrapped to avoid effects from periodic boundary conditions. For each voxel, we tracked the occupancy of fibers of type j and time-averaged over blocks of consecutive frames to reduce temporal correlations. Voxels whose time-averaged total occupancy fell below a prescribed threshold (0.2) were classified as effectively empty and excluded from subsequent analysis, ensuring that only the dense phase contributed to the demixing metric.

For each voxel, we computed fluctuations in local composition relative to the global mean. These fluctuations were decomposed into components parallel and perpendicular to the compositional contrast between the two fiber species, corresponding to composition and total-density fluctuations. The demixing index D was defined as the ratio of the variance of compositional fluctuations to that of density fluctuations. By construction, larger values of D indicate stronger local demixing, whereas smaller values correspond to more homogeneous mixing.

To facilitate comparison with experimental colocalization measurements, we converted the demixing index to a mixing score, defined as 1−D, so that larger values correspond to greater mixing. All reported values were averaged over independent simulation replicas.

## SUPPLEMENTARY FIGURES

**Fig. S1:**
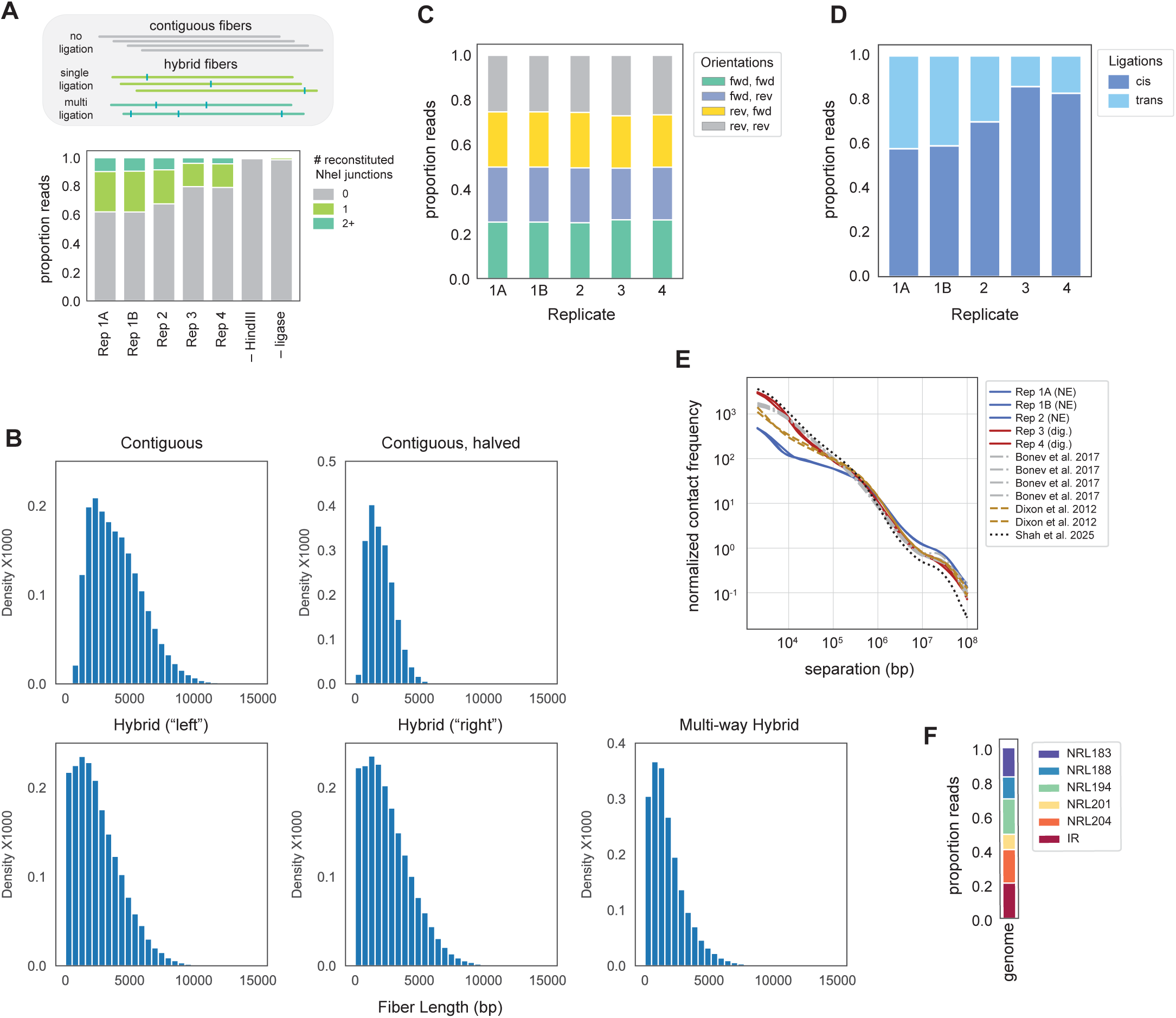
Key statistics for LASSI fibers, including proportion of ligation junctions, pair directions, cis/trans reads, and contact frequency. **A.)** <u>Top</u>: Schematic of contiguous and hybrid fibers found by LASSI. Ligation junctions, identified by the NheI restriction enzyme recognition site, shown in teal. <u>Bottom</u>: Proportion of reads with ligation junctions (0, 1, or 2+) from samples that underwent the full LASSI protocol (labeled as replicates) compared to controls without restriction enzyme or without ligase. For Replicates 1 and 2, nuclei were extracted as previously described in the SAMOSA protocol. For Replicates 3 and 4, cells were treated with digitonin instead of nuclear extraction buffer. Replicates 1A and 1B are technical replicates, while all others are biological replicates. **B.)** Length distributions for each type of molecule (contiguous, contiguous divided in half, and single or multi-way hybrids). **C.)** Proportion of strand direction combinations in each replicate. **D.)** Proportion of ligations in *cis* (on the same chromosome) or in *trans* (on different chromosomes) across each replicate. **E.)** Comparison of contact frequency between LASSI with either nuclear extraction or digitonin permeabilization, compared to 4-cutter in situ(*4*), 6-cutter dilution(*48*), and kit-based Hi-C(*47*). **F.)** Genome-wide proportion of each fiber type

**Fig. S2:**
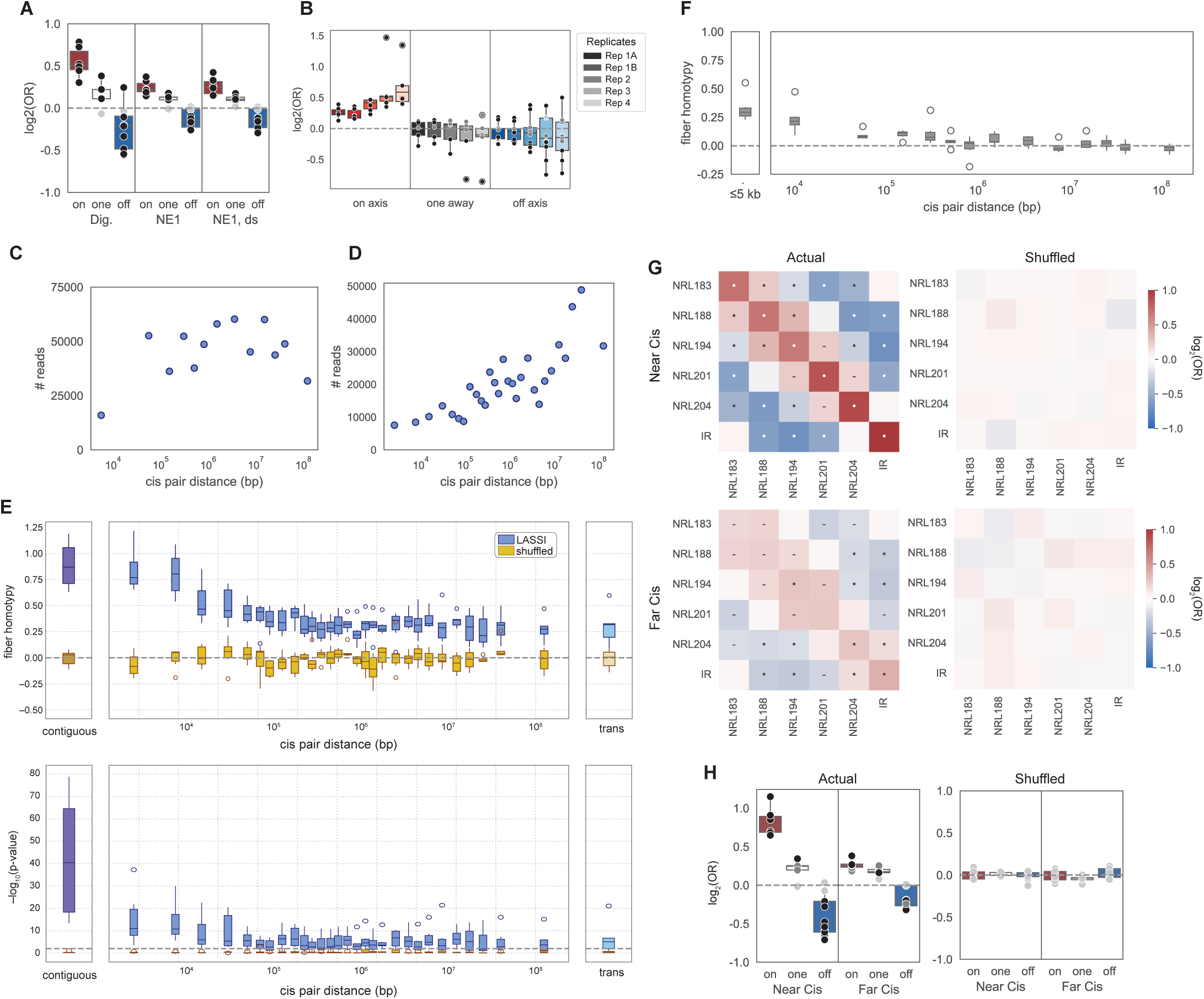
Read counts and alternative sampling for LASSI across cis pair distances. **A.)** Boxplot representations for different nuclei preparation methods (Dig. for digitonin; NE1 for nuclear extraction; NE1, ds to indicate nuclear extraction downsampled to the same number of hybrid fibers as digitonin). **B.)** Boxplot representations separated by replicate. **C.)** Read count for cis pair distance bins in Fig. 2. Bins were selected for best read depth and distribution by cis pair distance. Contiguous and trans pairs were downsampled to 50,000 reads. **D.)** Read count for bins in Fig. S2. Bins were selected for best read depth and distribution by cis pair distance. Contiguous and trans pairs were downsampled to 50,000 reads. **E.)** Top: Fiber homotypy scores for finer-grained bins, where blue bars show data from LASSI fibers and yellow bars show shuffled controls per each bin. Bottom: Storey’s *q* value for fiber homotypy scores shown in top panel. **F.)** Fiber homotypy scores calculated for random pairs from genome, binned by genomic distance between each “side” of the random pair. Each bin contains 50,000 pairs. **G.)** Left: Heatmaps depictions for near cis (≤10 kb pair distance) or far cis (> 60 Mb pair distance) hybrid fibers. Right: The same pool of “sides” that contributed to the hybrid fibers in left heatmaps but shuffled to remove spatial proximity correspondence. **H.)** Boxplot representation of heatmaps from S2D. Each dot is a fiber type. Black dots indicate q-value ≤ 0.01, dark grey dots indicate q-value ≤ 0.05, and light grey dots indicate q-value > 0.05.

**Fig. S3:**
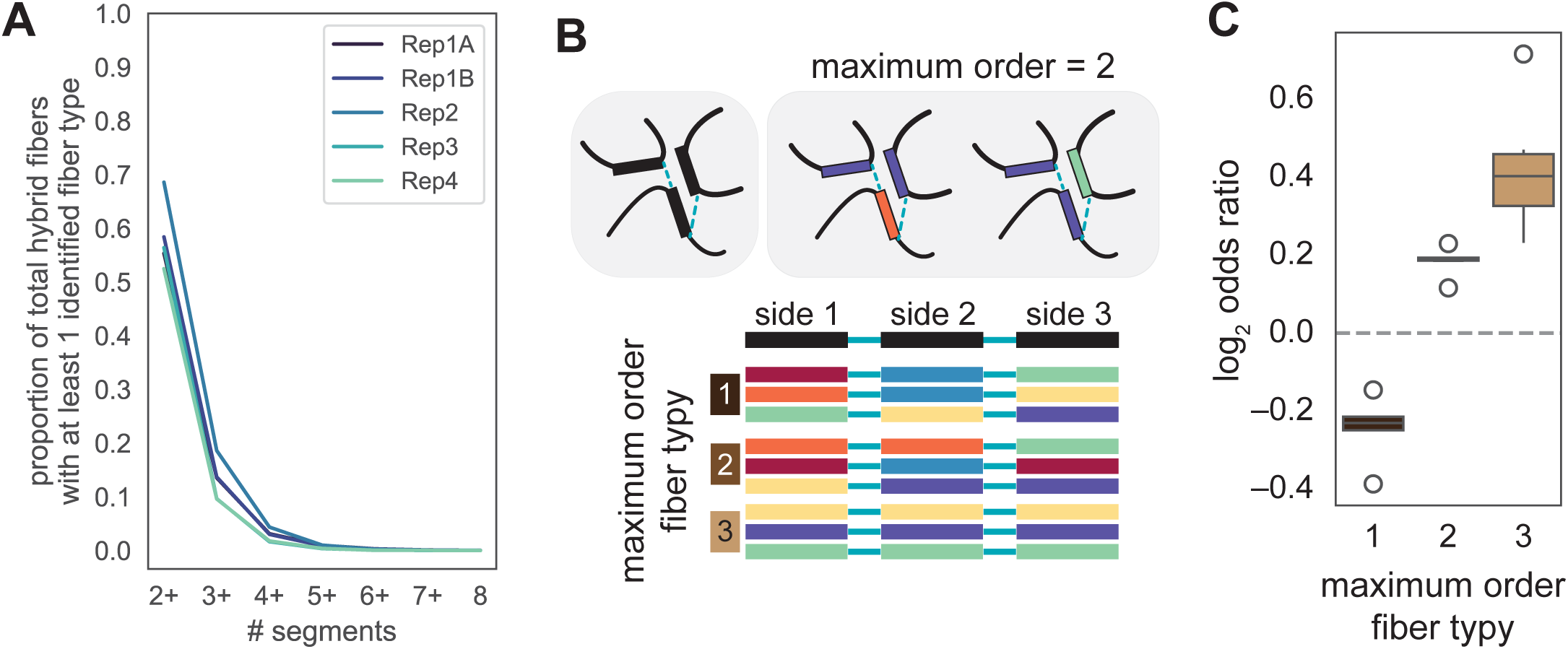
LASSI fibers that consist of multiple (>2) junctions show higher order homotypy. **A.)** Proportion of multi-way reads with at least 2 or more segments per hybrid fiber across each replicate. **B.)** Top, left: Example of 3-way hybrid fiber. Top, right: Examples in which the maximum order of homotypy is pairwise. Bottom: Examples of each order of homotypy as indicated, where fiber types are represented by color. **C.)** Log_2_ odds ratio of mutually exclusive, exactly paired, or higher-order shared fiber types for each three-segment hybrid fiber.

**Fig. S4:**
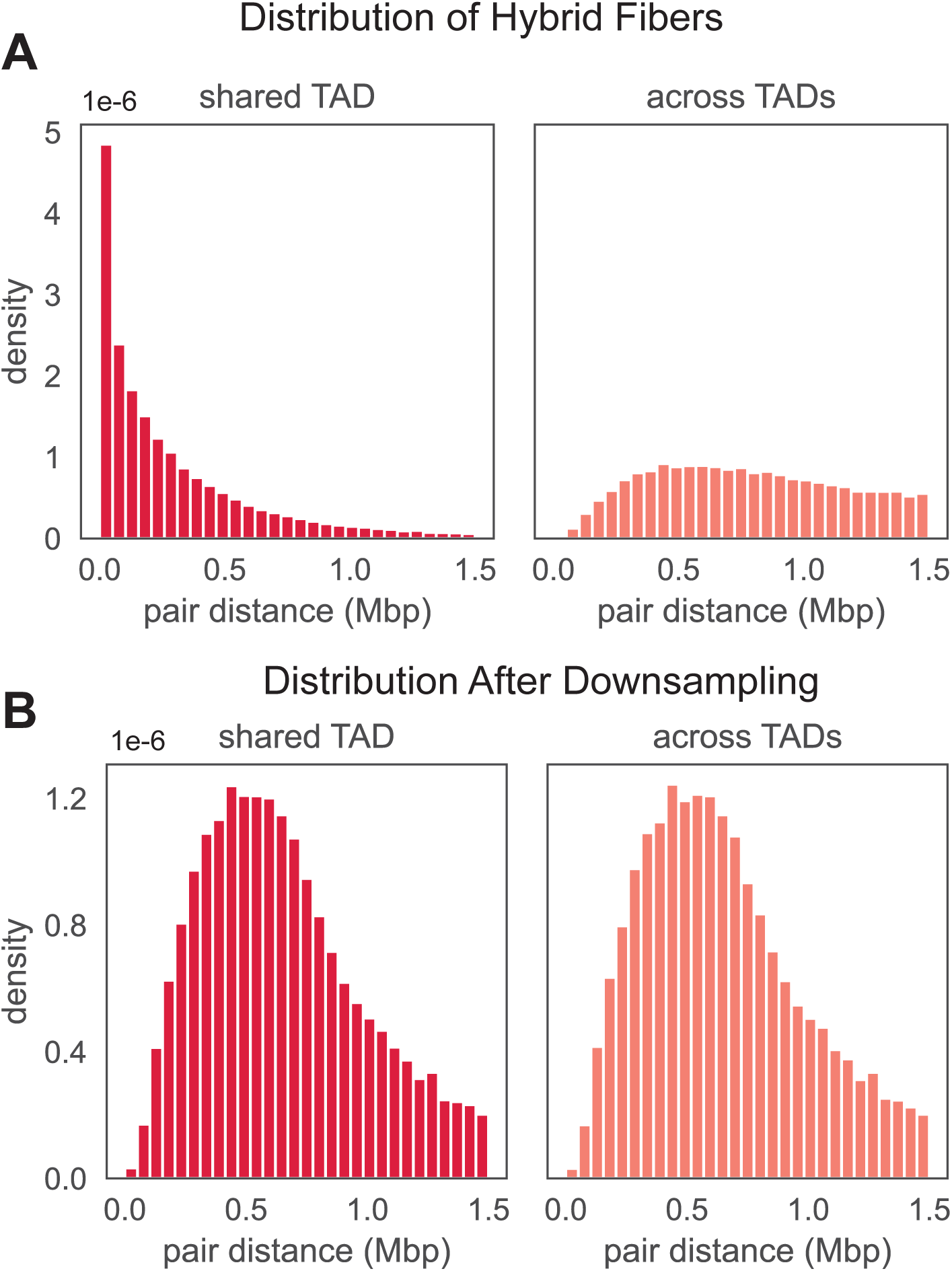
Downsampling reads to match cis pair distances for comparing within and across TAD hybrid fibers. **A.)** Read distribution by pair distance of hybrid fibers where each side is within the same TAD (“shared TAD”) or different TADs (“across TAD”). Hybrid fibers within shared TADs tend to also be close together in cis distance. **B.)** Read distribution by pair distance after downsampling to the minimum between shared or across TADs. Read distributions now match.

**Fig. S5:**
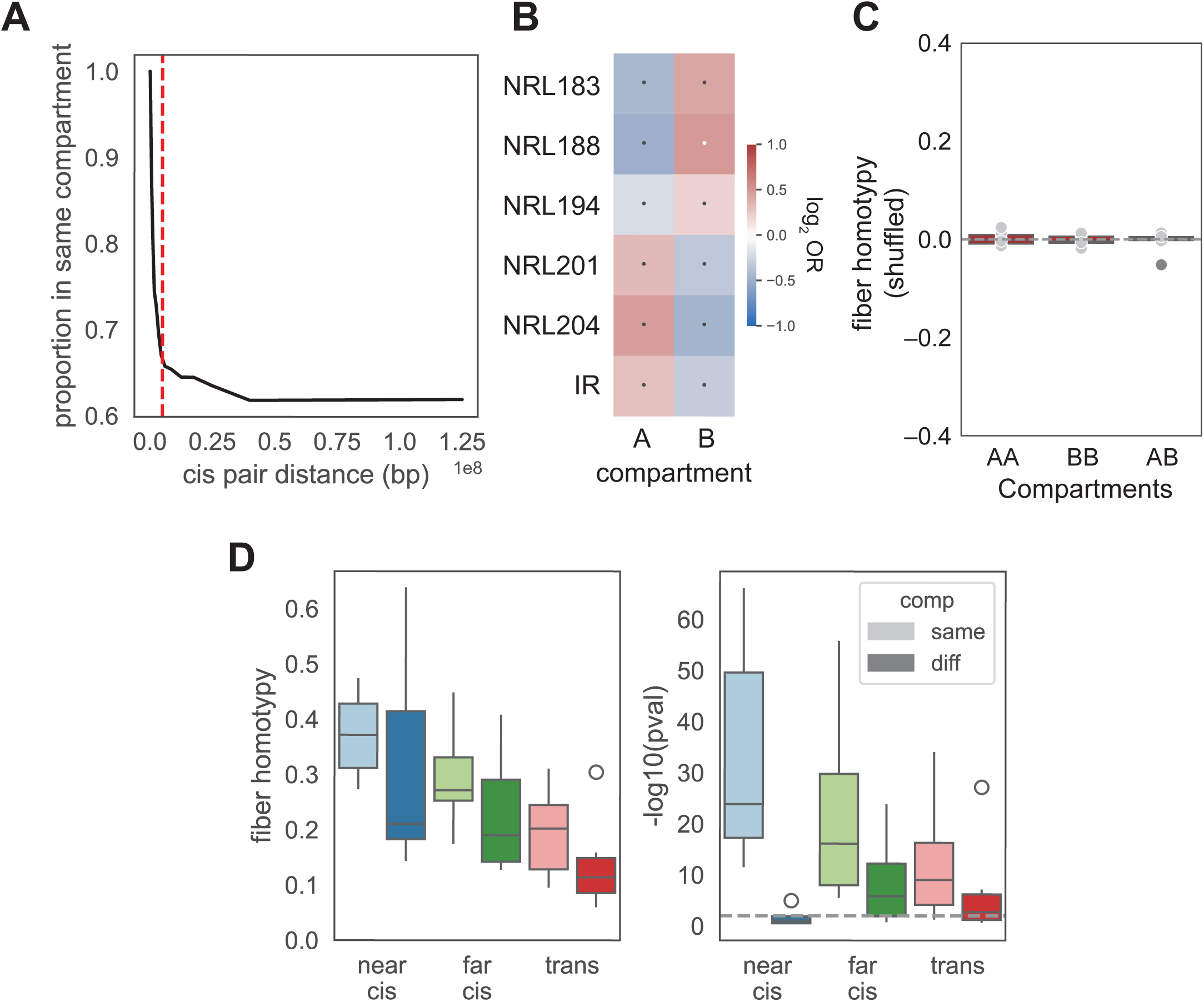
Key statistics and comparisons for fiber homotypy in A/B compartments. **A.)** Proportion of hybrid fibers in the same compartment by cis pair distance. Dashed red line indicates 5 Mb pair distance. **B.)** Fiber type enrichments by compartment. **C.)** Homotypy of shuffled pairs of hybrid fibers with either shared compartments (A compartment, B compartment) or across compartments. Each dot is a fiber type. Dark grey dots indicate p-value ≤ 0.05, and light grey dots indicate p-value > 0.05. **D.)** Fiber homotypy and associated *q* values within or across compartments at near cis (≤ 1 Mb), far cis (> 1 Mb), or trans.

**Fig. S6:**
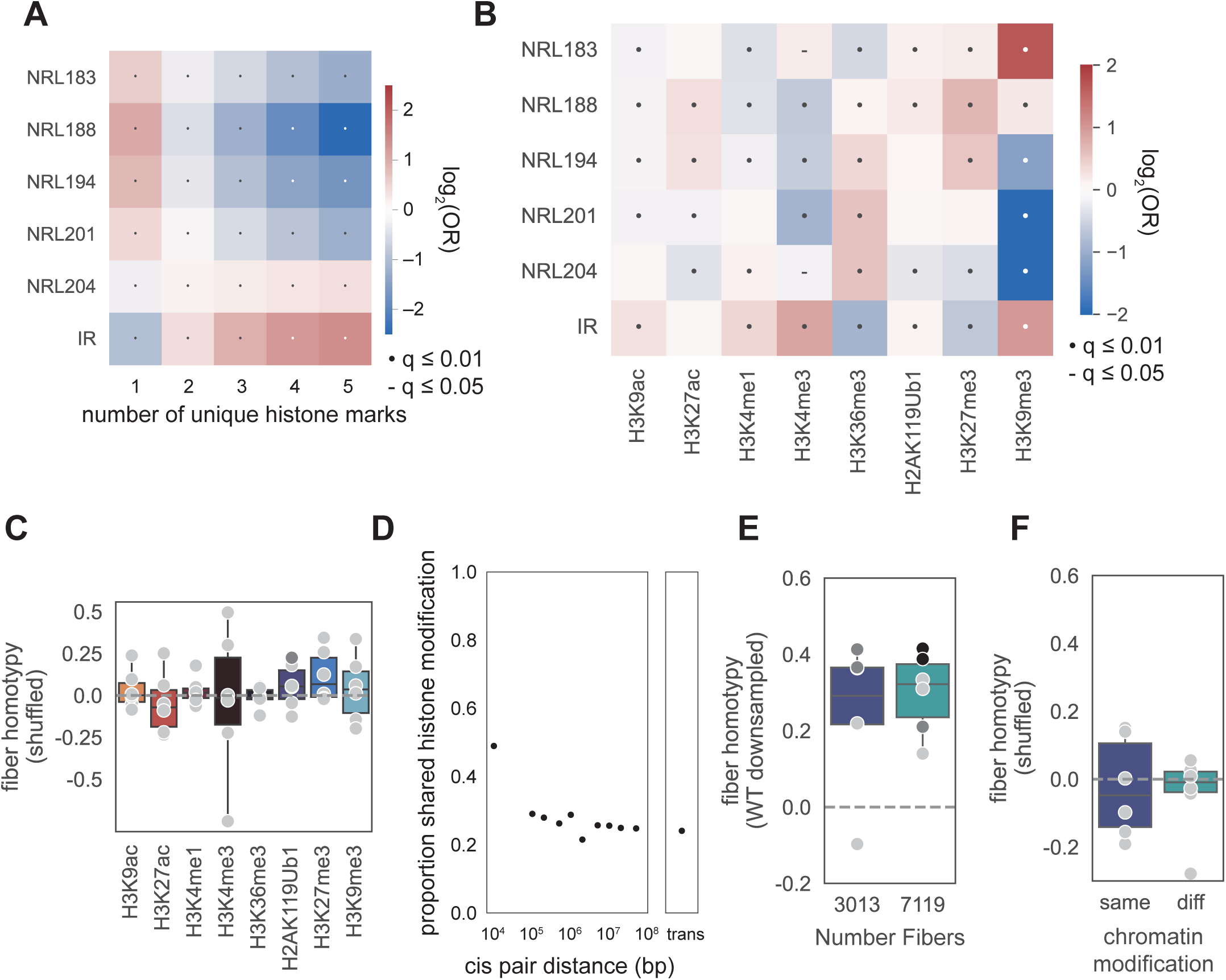
Histone modifications influence fiber type, yet increased fiber homotypy is not due to differential distributions of fiber type. **A.)** Fiber type enrichments by number of unique histone marks (as denoted by ENCODE ChIP-seq peaks) per each side per hybrid fiber. Any hybrid fibers with more than 1 unique histone mark were filtered out for subsequent analyses. **B.)** Fiber type enrichments by histone modification per each side per hybrid fiber. **C.)** Fiber homotypy in shuffled hybrid fibers where both sides have the same histone modification. Each dot is a fiber type. Dark grey dots indicate p-value ≤ 0.05, and light grey dots indicate p-value > 0.05. **D.)** Proportion of hybrid fibers that contain ENCODE ChIP peaks with shared histone modifications by cis pair distance. **E.)** Fiber homotypy calculated from downsampled WT hybrid fibers without filtering by histone modification, corresponding to the number of fibers with the same or different histone modifications. Each dot is a fiber type. Black dots indicate p-value ≤ 0.01, dark grey dots indicate p-value ≤ 0.05, and light grey dots indicate p-value > 0.05. **F.)** Fiber homotypy for shuffled hybrid fibers with the same (left, N=3013) or different (right, N=7119) chromatin modifications. Each dot is a fiber type. Light grey dots indicate p-value > 0.05.

**Fig. S7:**
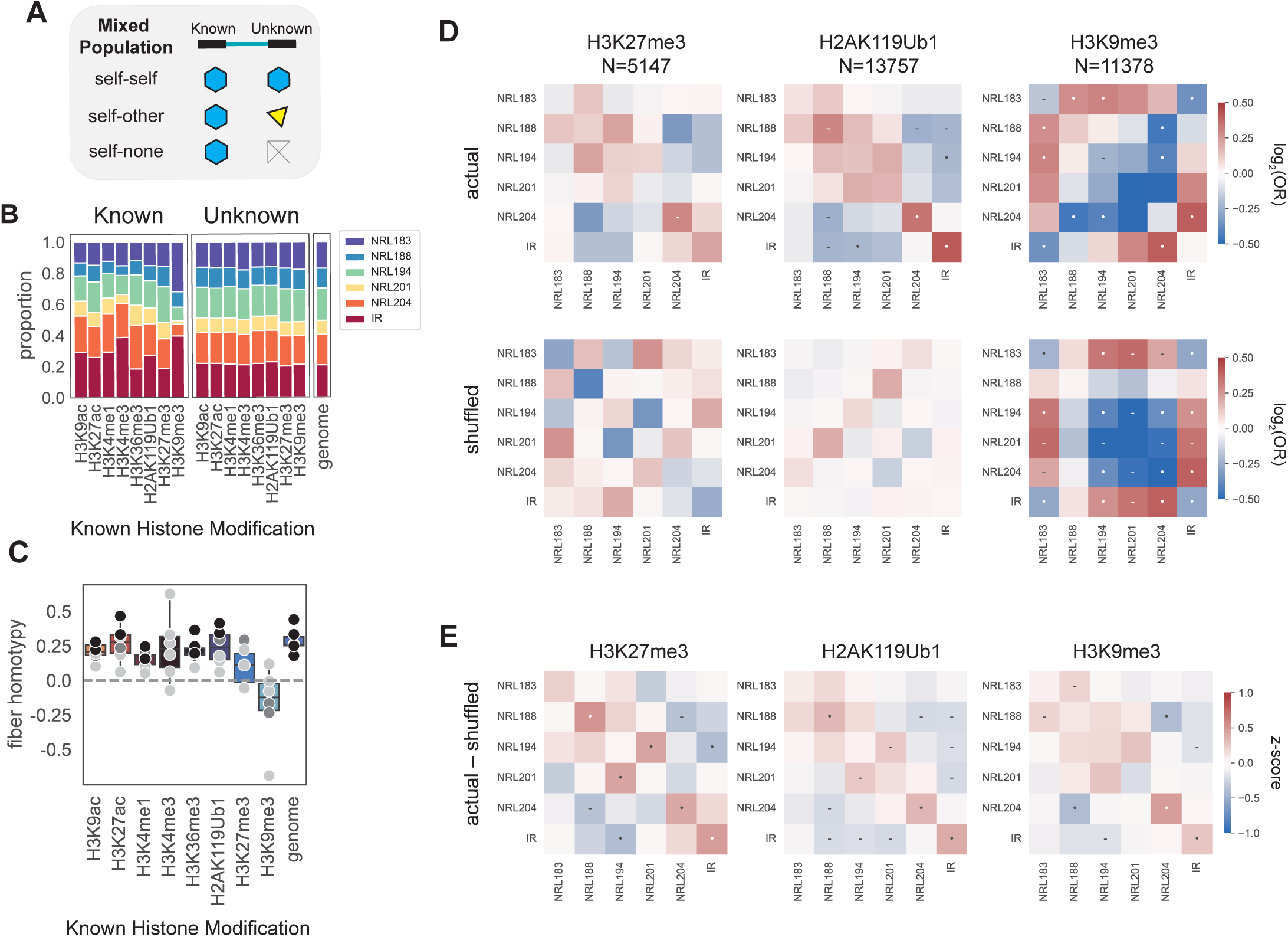
Histone modification alone does not inform nucleosome spacing in trans. **A.)** Schematic of all potential combinations where a histone modification is identified on only one side of a hybrid fiber. **B.)** Differential fiber type enrichments in fibers with an identified histone modification in cis (“known” side”) vs. in trans (“unknown” side). **C.)** Fiber homotypy in hybrid fibers with one-sided histone modifications. . Each dot is a fiber type. Black dots indicate p-value ≤ 0.01, dark grey dots indicate p-value ≤ 0.05, and light grey dots indicate p-value > 0.05. **D.)** Actual (top) vs. shuffled (bottom) log_2_(OR) heatmaps of fiber type pair enrichments in either facultative heterochromatin (H3K27me3 and H2AK119Ub1) or constitutive heterochromatin (H3K9me3). **E.)** Z-scores of actual vs. shuffled calculated from S7D.

**Fig. S8:**
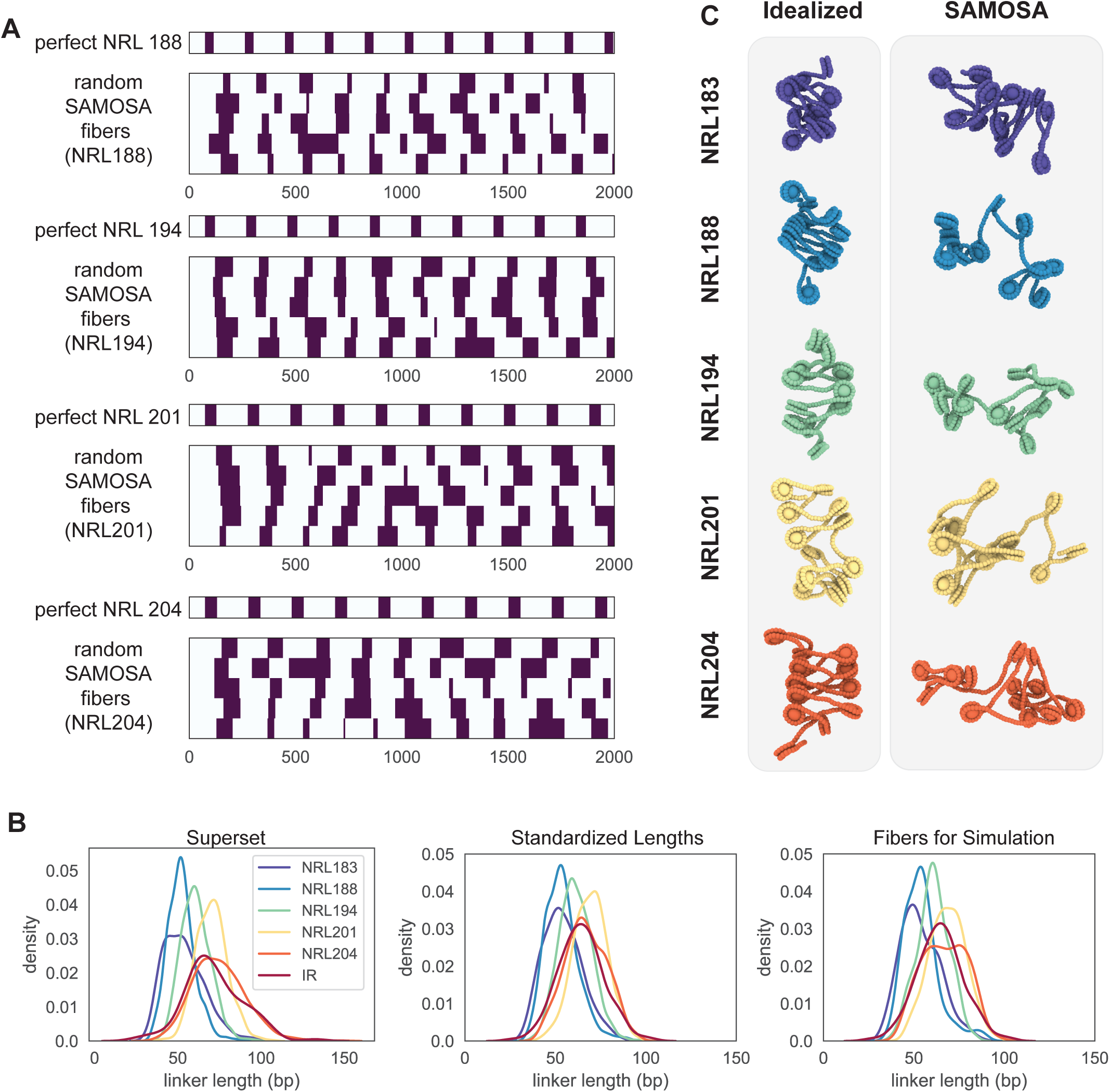
Comparison of chromatin arrays with idealized nucleosome repeat lengths and in vivo as footprinted by SAMOSA. **A.)** Comparison of footprints between a canonical array (top, per panel) with example LASSI footprinting data from each regular fiber type. All arrays begin at the center of the first nucleosome. **B.)** Mean linker in all SAMOSA data (“Superset”) compared to filtered (“Standardized Lengths”) and subsampled fibers used in simulations. **C.)** Simulations of canonical arrays with exact nucleosome spacings compared to simulations of one sampled SAMOSA-informed fiber.

**Fig. S9:**
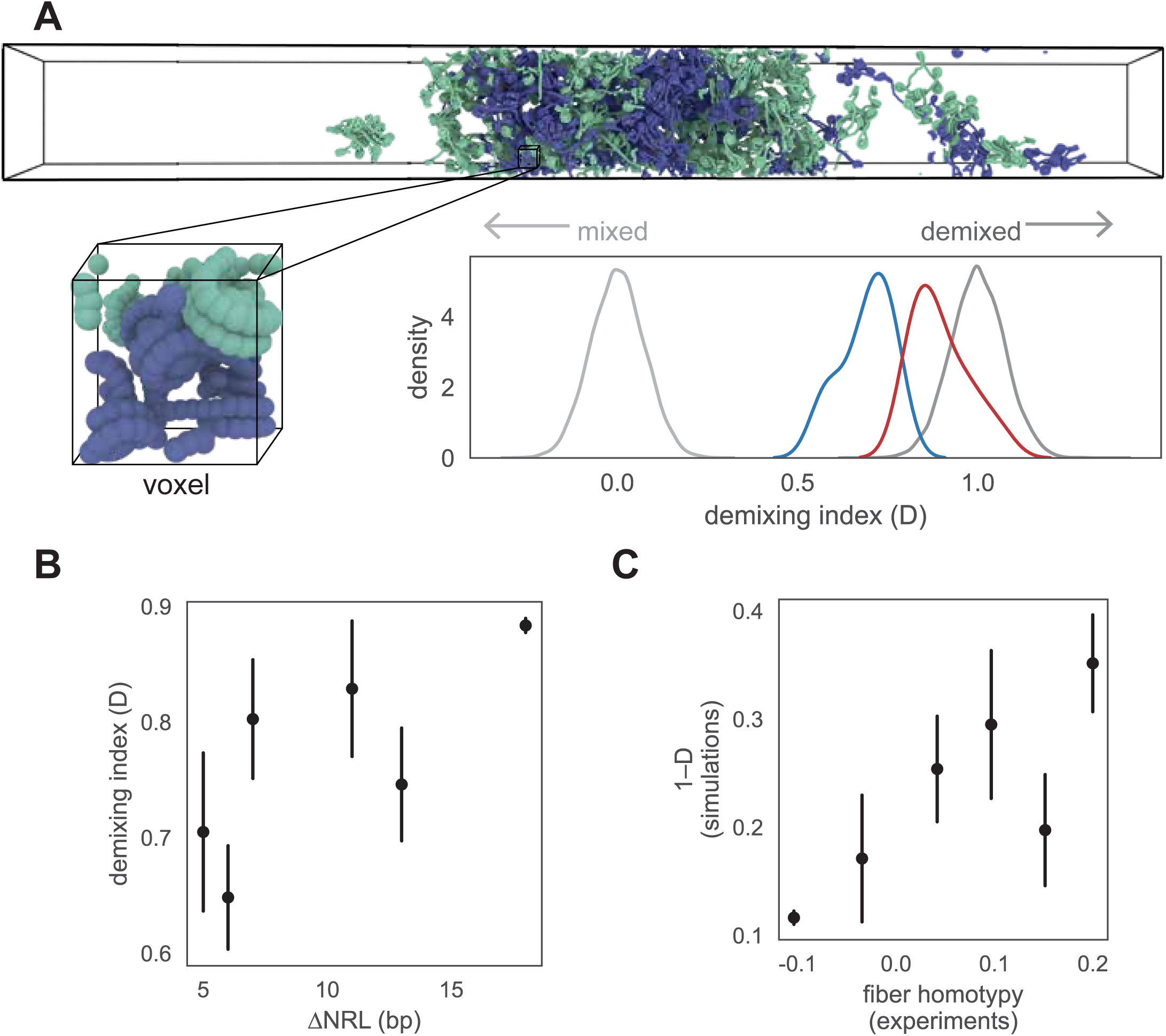
Voxel analysis on SAMOSA-informed simulations. **A.)** Schematic for analysis, including full slab simulations (top) and an example voxel (cutout). From these voxels (see Methods), a demixing index was calculated, where D=0 perfect mixing and D=1 represents perfect demixing, with representative values in gray. Values from actual simulations from either the upper half (generally demixed, red) or lower half (generally mixed, blue) are overlayed. **B.)** Correlation of fiber type NRL differences with demixing index (R=0.56, p=0.016). Interactions with NRL204 are omitted. Error bars show ± 1 SEM. **C.)** Correlation (R=0.61, p=0.008) of 1–D (mixing) with experimental fiber heterotypy. Interactions with NRL204 are omitted. Error bars show ± 1 SEM.

**Fig. S10:**
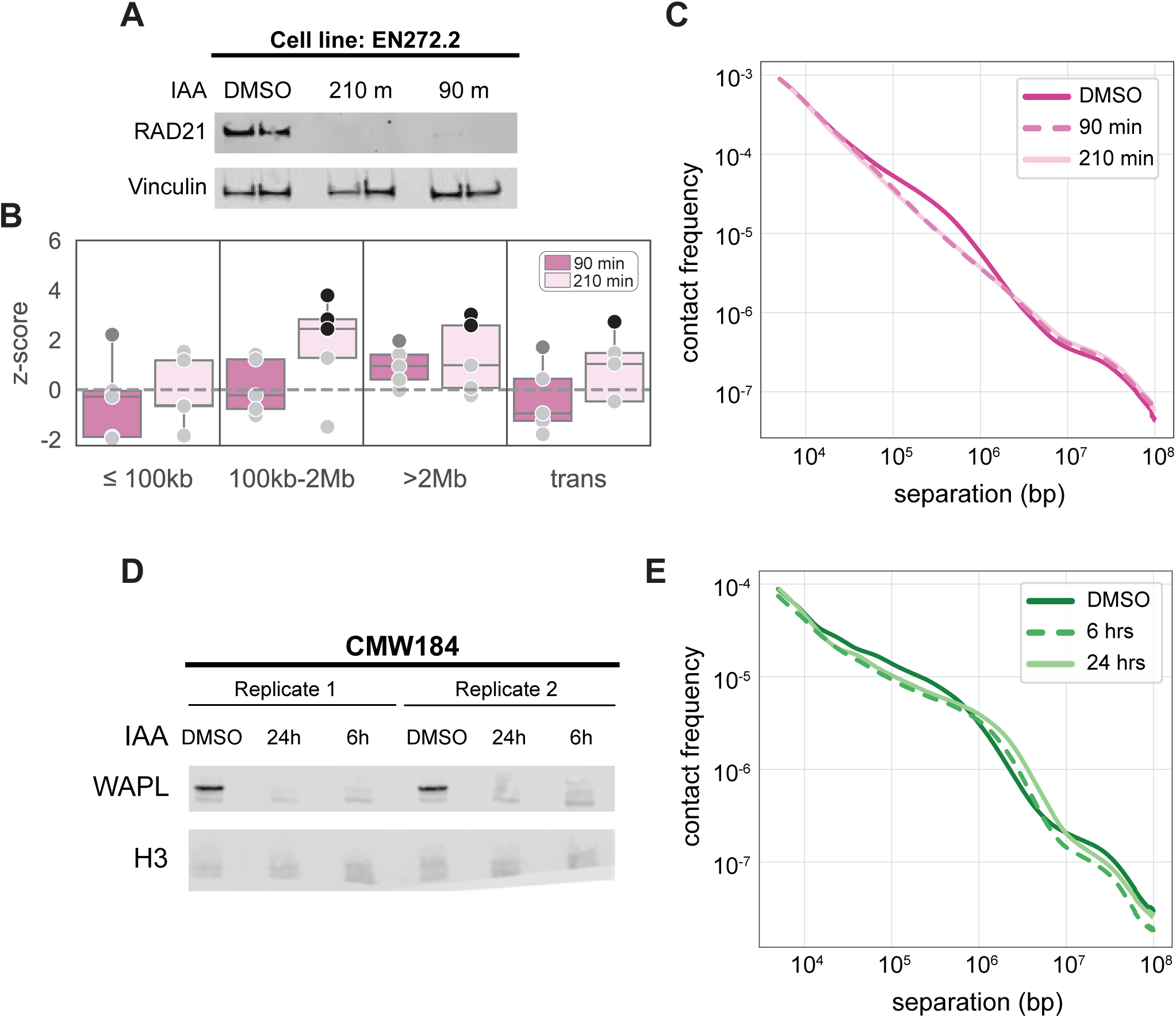
RAD21 and WAPL degron validation and additional characterization. **A.)** Western blot of knockdown for RAD21 in cell line EN272.2. **B.)** Z-score of each depletion when compared to DMSO condition separated by cis pair distance. Each dot is a fiber type. Black dots indicate p-value ≤ 0.01, dark grey dots indicate p-value ≤ 0.05, and light grey dots indicate p-value > 0.05. **C.)** Contact frequency for RAD21 depletion experiment as measured by LASSI. **D.)** Western blot of knockdown for WAPL in cell line CMW184. **E.)** Contact frequency for WAPL depletion experiment as measured by LASSI.

## REFERENCES

1. T. Misteli, The Self-Organizing Genome: Principles of Genome Architecture and Function. Cell 183, 28–45 (2020).

2. M. J. Rowley, V. G. Corces, Organizational principles of 3D genome architecture. Nat. Rev. Genet. 19, 789–800 (2018).

3. C. R. Clapier, J. Iwasa, B. R. Cairns, C. L. Peterson, Mechanisms of action and regulation of ATP-dependent chromatin-remodelling complexes. Nat. Rev. Mol. Cell Biol. 18, 407–422 (2017).

4. B. Bonev, N. Mendelson Cohen, Q. Szabo, L. Fritsch, G. L. Papadopoulos, Y. Lubling, X. Xu, X. Lv, J.-P. Hugnot, A. Tanay, G. Cavalli, Multiscale 3D Genome Rewiring during Mouse Neural Development. Cell 171, 557–572.e24 (2017).

5. S. Ide, S. Tamura, K. Maeshima, Chromatin behavior in living cells: Lessons from single-nucleosome imaging and tracking. BioEssays 44, 2200043 (2022).

6. A. Routh, S. Sandin, D. Rhodes, Nucleosome repeat length and linker histone stoichiometry determine chromatin fiber structure. Proc. Natl. Acad. Sci. 105, 8872–8877 (2008).

7. P. J. J. Robinson, L. Fairall, V. A. T. Huynh, D. Rhodes, EM measurements define the dimensions of the “30-nm” chromatin fiber: Evidence for a compact, interdigitated structure. Proc. Natl. Acad. Sci. 103, 6506–6511 (2006).

8. H. D. Ou, S. Phan, T. J. Deerinck, A. Thor, M. H. Ellisman, C. C. O’Shea, ChromEMT: Visualizing 3D chromatin structure and compaction in interphase and mitotic cells. Science 357, eaag0025 (2017).

9. H. Zhou, J. Huertas, M. J. Maristany, K. Russell, J. H. Hwang, R.-W. Yao, N. Samanta, J. Hutchings, R. Billur, M. Shiozaki, X. Zhao, L. K. Doolittle, B. A. Gibson, A. Soranno, M. Riggi, J. R. Espinosa, Z. Yu, E. Villa, R. Collepardo-Guevara, M. K. Rosen, Multiscale structure of chromatin condensates explains phase separation and material properties. Science 390, eadv6588 (2025).

10. B. A. Gibson, L. K. Doolittle, M. W. G. Schneider, L. E. Jensen, N. Gamarra, L. Henry, D. W. Gerlich, S. Redding, M. K. Rosen, Organization of Chromatin by Intrinsic and Regulated Phase Separation. Cell 179, 470–484.e21 (2019).

11. B. A. Gibson, C. Blaukopf, T. Lou, L. Chen, L. K. Doolittle, I. Finkelstein, G. J. Narlikar, D. W. Gerlich, M. K. Rosen, In diverse conditions, intrinsic chromatin condensates have liquid-like material properties. Proc. Natl. Acad. Sci. 120, e2218085120 (2023).

12. L. Chen, M. J. Maristany, S. E. Farr, J. Luo, B. A. Gibson, L. K. Doolittle, J. R. Espinosa, J. Huertas, S. Redding, R. Collepardo-Guevara, M. K. Rosen, Nucleosome spacing can fine-tune higher-order chromatin assembly. Nat. Commun. 16, 6315 (2025).

13. M. Falk, Y. Feodorova, N. Naumova, M. Imakaev, B. R. Lajoie, H. Leonhardt, B. Joffe, J. Dekker, G. Fudenberg, I. Solovei, L. A. Mirny, Heterochromatin drives compartmentalization of inverted and conventional nuclei. Nature 570, 395–399 (2019).

14. D. Hnisz, K. Shrinivas, R. A. Young, A. K. Chakraborty, P. A. Sharp, A Phase Separation Model for Transcriptional Control. Cell 169, 13–23 (2017).

15. N. J. Abdulhay, C. P. McNally, L. J. Hsieh, S. Kasinathan, A. Keith, L. S. Estes, M. Karimzadeh, J. G. Underwood, H. Goodarzi, G. J. Narlikar, V. Ramani, Massively multiplex single-molecule oligonucleosome footprinting. eLife 9, e59404 (2020).

16. N. J. Abdulhay, L. J. Hsieh, C. P. McNally, M. S. Ostrowski, C. M. Moore, M. Ketavarapu, S. Kasinathan, A. S. Nanda, K. Wu, U. S. Chio, Z. Zhou, H. Goodarzi, G. J. Narlikar, V. Ramani, Nucleosome density shapes kilobase-scale regulation by a mammalian chromatin remodeler. Nat. Struct. Mol. Biol. 30, 1571–1581 (2023).

17. A. B. Stergachis, B. M. Debo, E. Haugen, L. S. Churchman, J. A. Stamatoyannopoulos, Single-molecule regulatory architectures captured by chromatin fiber sequencing. Science 368, 1449–1454 (2020).

18. A. R. Krebs, D. Imanci, L. Hoerner, D. Gaidatzis, L. Burger, D. Schübeler, Genome-wide Single-Molecule Footprinting Reveals High RNA Polymerase II Turnover at Paused Promoters. Mol. Cell 67, 411–422.e4 (2017).

19. T. Nagano, Y. Lubling, T. J. Stevens, S. Schoenfelder, E. Yaffe, W. Dean, E. D. Laue, A. Tanay, P. Fraser, Single-cell Hi-C reveals cell-to-cell variability in chromosome structure. Nature 502, 59–64 (2013).

20. B. Bintu, L. J. Mateo, J.-H. Su, N. A. Sinnott-Armstrong, M. Parker, S. Kinrot, K. Yamaya, A. N. Boettiger, X. Zhuang, Super-resolution chromatin tracing reveals domains and cooperative interactions in single cells. Science 362, eaau1783 (2018).

21. M. A. Ricci, C. Manzo, M. F. García-Parajo, M. Lakadamyali, M. P. Cosma, Chromatin Fibers Are Formed by Heterogeneous Groups of Nucleosomes In Vivo. Cell 160, 1145–1158 (2015).

22. T. Nozaki, R. Imai, M. Tanbo, R. Nagashima, S. Tamura, T. Tani, Y. Joti, M. Tomita, K. Hibino, M. T. Kanemaki, K. S. Wendt, Y. Okada, T. Nagai, K. Maeshima, Dynamic Organization of Chromatin Domains Revealed by Super-Resolution Live-Cell Imaging. Mol. Cell 67, 282–293.e7 (2017).

23. L. J. Mateo, S. E. Murphy, A. Hafner, I. S. Cinquini, C. A. Walker, A. N. Boettiger, Visualizing DNA folding and RNA in embryos at single-cell resolution. Nature 568, 49–54 (2019).

24. G. Fudenberg, M. Imakaev, C. Lu, A. Goloborodko, N. Abdennur, L. A. Mirny, Formation of Chromosomal Domains by Loop Extrusion. Cell Rep. 15, 2038–2049 (2016).

25. S. S. P. Rao, M. H. Huntley, N. C. Durand, E. K. Stamenova, I. D. Bochkov, J. T. Robinson, A. L. Sanborn, I. Machol, A. D. Omer, E. S. Lander, E. L. Aiden, A 3D Map of the Human Genome at Kilobase Resolution Reveals Principles of Chromatin Looping. Cell 159, 1665–1680 (2014).

26. J. Dekker, K. Rippe, M. Dekker, N. Kleckner, Capturing Chromosome Conformation. Science 295, 1306–1311 (2002).

27. E. Lieberman-Aiden, N. L. van Berkum, L. Williams, M. Imakaev, T. Ragoczy, A. Telling, I. Amit, B. R. Lajoie, P. J. Sabo, M. O. Dorschner, R. Sandstrom, B. Bernstein, M. A. Bender, M. Groudine, A. Gnirke, J. Stamatoyannopoulos, L. A. Mirny, E. S. Lander, J. Dekker, Comprehensive Mapping of Long-Range Interactions Reveals Folding Principles of the Human Genome. Science 326, 289–293 (2009).

28. B. Akgol Oksuz, L. Yang, S. Abraham, S. V. Venev, N. Krietenstein, K. M. Parsi, H. Ozadam, M. E. Oomen, A. Nand, H. Mao, R. M. J. Genga, R. Maehr, O. J. Rando, L. A. Mirny, J. H. Gibcus, J. Dekker, Systematic evaluation of chromosome conformation capture assays. Nat. Methods 18, 1046–1055 (2021).

29. A. S. Nanda, K. Wu, I. Irkliyenko, B. Woo, M. S. Ostrowski, A. S. Clugston, L. C. Sayles, L. Xu, A. T. Satpathy, H. G. Nguyen, E. Alejandro Sweet-Cordero, H. Goodarzi, S. Kasinathan, V. Ramani, Direct transposition of native DNA for sensitive multimodal single-molecule sequencing. Nat. Genet. 56, 1300–1309 (2024).

30. C. Moore, E. Wong, U. Kaur, U. S. Chio, Z. Zhou, M. Ostrowski, K. Wu, I. Irkliyenko, S. Wang, V. Ramani, G. J. Narlikar, ATP-dependent remodeling of chromatin condensates reveals distinct mesoscale outcomes. Science 390, eadr0018 (2025).

31. M. S. Ostrowski, M. G. Yang, C. P. McNally, N. J. Abdulhay, S. Wang, K. Renduchintala, I. Irkliyenko, A. Biran, B. T. L. Chew, A. D. Midha, E. V. Wong, J. Sandoval, I. H. Jain, A. Groth, E. P. Nora, H. Goodarzi, V. Ramani, The single-molecule accessibility landscape of newly replicated mammalian chromatin. Cell 188, 237–252.e19 (2025).

32. M. G. Yang, H. J. Richter, S. Wang, C. P. McNally, N. Harris, S. Dhillon, M. Maresca, E. de Wit, H. Willenbring, J. Maher, H. Goodarzi, V. Ramani, Pervasive and programmed nucleosome distortion patterns on single mammalian chromatin fibers. bioRxiv [Preprint] (2025). 10.1101/2025.01.17.633622.

33. Z. Shipony, G. K. Marinov, M. P. Swaffer, N. A. Sinnott-Armstrong, J. M. Skotheim, A. Kundaje, W. J. Greenleaf, Long-range single-molecule mapping of chromatin accessibility in eukaryotes. Nat. Methods 17, 319–327 (2020).

34. S. Battaglia, K. Dong, J. Wu, Z. Chen, F. J. Najm, Y. Zhang, M. M. Moore, V. Hecht, N. Shoresh, B. E. Bernstein, Long-range phasing of dynamic, tissue-specific and allele-specific regulatory elements. Nat. Genet. 54, 1504–1513 (2022).

35. N. Altemose, A. Maslan, O. K. Smith, K. Sundararajan, R. R. Brown, R. Mishra, A. M. Detweiler, N. Neff, K. H. Miga, A. F. Straight, A. Streets, DiMeLo-seq: a long-read, single-molecule method for mapping protein–DNA interactions genome wide. Nat. Methods 19, 711–723 (2022).

36. I. Lee, R. Razaghi, T. Gilpatrick, M. Molnar, A. Gershman, N. Sadowski, F. J. Sedlazeck, K. D. Hansen, J. T. Simpson, W. Timp, Simultaneous profiling of chromatin accessibility and methylation on human cell lines with nanopore sequencing. Nat. Methods 17, 1191–1199 (2020).

37. T. K. Kelly, Y. Liu, F. D. Lay, G. Liang, B. P. Berman, P. A. Jones, Genome-wide mapping of nucleosome positioning and DNA methylation within individual DNA molecules. Genome Res. 22, 2497–2506 (2012).

38. Y. Xie, F. Ruan, Y. Li, M. Luo, C. Zhang, Z. Chen, Z. Xie, Z. Weng, W. Chen, W. Chen, Y. Fang, Y. Sun, M. Guo, J. Wang, S. Xu, H. Wang, C. Tang, Spatial chromatin accessibility sequencing resolves high-order spatial interactions of epigenomic markers. eLife 12, RP87868 (2024).

39. F. Noack, S. Vangelisti, N. Ditzer, F. Chong, M. Albert, B. Bonev, Joint epigenome profiling reveals cell-type-specific gene regulatory programmes in human cortical organoids. Nat. Cell Biol. 25, 1873–1883 (2023).

40. A. S. Deshpande, N. Ulahannan, M. Pendleton, X. Dai, L. Ly, J. M. Behr, S. Schwenk, W. Liao, M. A. Augello, C. Tyer, P. Rughani, S. Kudman, H. Tian, H. G. Otis, E. Adney, D. Wilkes, J. M. Mosquera, C. E. Barbieri, A. Melnick, D. Stoddart, D. J. Turner, S. Juul, E. Harrington, M. Imieliński, Identifying synergistic high-order 3D chromatin conformations from genome-scale nanopore concatemer sequencing. Nat. Biotechnol. 40, 1488–1499 (2022).

41. S. P. McGinty, G. Kaya, S. B. Sim, A. Makunin, R. L. Corpuz, M. A. Quail, M. Abuelanin, M. K. N. Lawniczak, S. M. Geib, J. Korlach, M. Y. Dennis, CiFi: accurate long-read chromosome conformation capture with low-input requirements. Nat. Commun. 17, 215 (2025).

42. V. Y. Goel, M. K. Huseyin, A. S. Hansen, Region Capture Micro-C reveals coalescence of enhancers and promoters into nested microcompartments. Nat. Genet. 55, 1048–1056 (2023).

43. J. C. Hamley, H. Li, N. Denny, D. Downes, J. O. J. Davies, Determining chromatin architecture with Micro Capture-C. Nat. Protoc. 18, 1687–1711 (2023).

44. H. Li, J. L. T. Dalgleish, G. Lister, M. J. Maristany, J. Huertas, A. M. Dopico-Fernandez, J. C. Hamley, N. Denny, G. Bloye, W. Zhang, L. Hentges, R. Doll, Y. Wei, M. Maresca, E. Dimitrova, L. Pytowski, E. A. J. Tunnacliffe, M. Kassouf, D. Higgs, E. de Wit, R. J. Klose, L. Schermelleh, R. Collepardo-Guevara, T. A. Milne, J. O. J. Davies, Mapping chromatin structure at base-pair resolution unveils a unified model of cis-regulatory element interactions. Cell 188, 7175–7193.e19 (2025).

45. P. Hua, M. Badat, L. L. P. Hanssen, L. D. Hentges, N. Crump, D. J. Downes, D. M. Jeziorska, A. M. Oudelaar, R. Schwessinger, S. Taylor, T. A. Milne, J. R. Hughes, D. R. Higgs, J. O. J. Davies, Defining genome architecture at base-pair resolution. Nature 595, 125–129 (2021).

46. T.-H. S. Hsieh, C. Cattoglio, E. Slobodyanyuk, A. S. Hansen, O. J. Rando, R. Tjian, X. Darzacq, Resolving the 3D Landscape of Transcription-Linked Mammalian Chromatin Folding. Mol. Cell 78, 539–553.e8 (2020).

47. R. Shah, M. M. C. Tortora, N. Louafi, H. Rahmaninejad, K. L. Hansen, E. C. Anderson, D. Wen, L. Giorgetti, G. Fudenberg, E. P. Nora, NIPBL dosage shapes genome folding by tuning the rate of cohesin loop extrusion. bioRxiv [Preprint] (2025). 10.1101/2025.08.14.667581.

48. J. R. Dixon, S. Selvaraj, F. Yue, A. Kim, Y. Li, Y. Shen, M. Hu, J. S. Liu, B. Ren, Topological domains in mammalian genomes identified by analysis of chromatin interactions. Nature 485, 376–380 (2012).

49. P. Olivares-Chauvet, Z. Mukamel, A. Lifshitz, O. Schwartzman, N. O. Elkayam, Y. Lubling, G. Deikus, R. P. Sebra, A. Tanay, Capturing pairwise and multi-way chromosomal conformations using chromosomal walks. Nature 540, 296–300 (2016).

50. W. Schwarzer, N. Abdennur, A. Goloborodko, A. Pekowska, G. Fudenberg, Y. Loe-Mie, N. A. Fonseca, W. Huber, C. H. Haering, L. Mirny, F. Spitz, Two independent modes of chromatin organization revealed by cohesin removal. Nature 551, 51–56 (2017).

51. E. P. Nora, A. Goloborodko, A.-L. Valton, J. H. Gibcus, A. Uebersohn, N. Abdennur, J. Dekker, L. A. Mirny, B. G. Bruneau, Targeted Degradation of CTCF Decouples Local Insulation of Chromosome Domains from Genomic Compartmentalization. Cell 169, 930–944.e22 (2017).

52. G. Spracklin, N. Abdennur, M. Imakaev, N. Chowdhury, S. Pradhan, L. A. Mirny, J. Dekker, Diverse silent chromatin states modulate genome compartmentalization and loop extrusion barriers. Nat. Struct. Mol. Biol. 30, 38–51 (2023).

53. F. Yue, Y. Cheng, A. Breschi, J. Vierstra, W. Wu, T. Ryba, R. Sandstrom, Z. Ma, C. Davis, B. D. Pope, Y. Shen, D. D. Pervouchine, S. Djebali, R. E. Thurman, R. Kaul, E. Rynes, A. Kirilusha, G. K. Marinov, B. A. Williams, D. Trout, H. Amrhein, K. Fisher-Aylor, I. Antoshechkin, G. DeSalvo, L.-H. See, M. Fastuca, J. Drenkow, C. Zaleski, A. Dobin, P. Prieto, J. Lagarde, G. Bussotti, A. Tanzer, O. Denas, K. Li, M. A. Bender, M. Zhang, R. Byron, M. T. Groudine, D. McCleary, L. Pham, Z. Ye, S. Kuan, L. Edsall, Y.-C. Wu, M. D. Rasmussen, M. S. Bansal, M. Kellis, C. A. Keller, C. S. Morrissey, T. Mishra, D. Jain, N. Dogan, R. S. Harris, P. Cayting, T. Kawli, A. P. Boyle, G. Euskirchen, A. Kundaje, S. Lin, Y. Lin, C. Jansen, V. S. Malladi, M. S. Cline, D. T. Erickson, V. M. Kirkup, K. Learned, C. A. Sloan, K. R. Rosenbloom, B. Lacerda de Sousa, K. Beal, M. Pignatelli, P. Flicek, J. Lian, T. Kahveci, D. Lee, W. James Kent, M. Ramalho Santos, J. Herrero, C. Notredame, A. Johnson, S. Vong, K. Lee, D. Bates, F. Neri, M. Diegel, T. Canfield, P. J. Sabo, M. S. Wilken, T. A. Reh, E. Giste, A. Shafer, T. Kutyavin, E. Haugen, D. Dunn, A. P. Reynolds, S. Neph, R. Humbert, R. Scott Hansen, M. De Bruijn, L. Selleri, A. Rudensky, S. Josefowicz, R. Samstein, E. E. Eichler, S. H. Orkin, D. Levasseur, T. Papayannopoulou, K.-H. Chang, A. Skoultchi, S. Gosh, C. Disteche, P. Treuting, Y. Wang, M. J. Weiss, G. A. Blobel, X. Cao, S. Zhong, T. Wang, P. J. Good, R. F. Lowdon, L. B. Adams, X.-Q. Zhou, M. J. Pazin, E. A. Feingold, B. Wold, J. Taylor, A. Mortazavi, S. M. Weissman, J. A. Stamatoyannopoulos, M. P. Snyder, R. Guigo, T. R. Gingeras, D. M. Gilbert, R. C. Hardison, M. A. Beer, B. Ren, A comparative encyclopedia of DNA elements in the mouse genome. Nature 515, 355–364 (2014).

54. S. E. Farr, E. J. Woods, J. A. Joseph, A. Garaizar, R. Collepardo-Guevara, Nucleosome plasticity is a critical element of chromatin liquid–liquid phase separation and multivalent nucleosome interactions. Nat. Commun. 12, 2883 (2021).

55. J. R. Espinosa, E. Sanz, C. Valeriani, C. Vega, On fluid-solid direct coexistence simulations: The pseudo-hard sphere model. J. Chem. Phys. 139, 144502 (2013).

56. W. Morton, R. Vacha, Quantifying (de)Mixing of Disordered Proteins in Molecular Dynamics Simulations.

57. S. S. P. Rao, S.-C. Huang, B. Glenn St Hilaire, J. M. Engreitz, E. M. Perez, K.-R. Kieffer-Kwon, A. L. Sanborn, S. E. Johnstone, G. D. Bascom, I. D. Bochkov, X. Huang, M. S. Shamim, J. Shin, D. Turner, Z. Ye, A. D. Omer, J. T. Robinson, T. Schlick, B. E. Bernstein, R. Casellas, E. S. Lander, E. L. Aiden, Cohesin Loss Eliminates All Loop Domains. Cell 171, 305–320.e24 (2017).

58. G. Wutz, C. Várnai, K. Nagasaka, D. A. Cisneros, R. R. Stocsits, W. Tang, S. Schoenfelder, G. Jessberger, M. Muhar, M. J. Hossain, N. Walther, B. Koch, M. Kueblbeck, J. Ellenberg, J. Zuber, P. Fraser, J. Peters, Topologically associating domains and chromatin loops depend on cohesin and are regulated by CTCF, WAPL, and PDS5 proteins. EMBO J. 36, 3573–3599 (2017).

59. J. H. I. Haarhuis, R. H. van der Weide, V. A. Blomen, J. O. Yáñez-Cuna, M. Amendola, M. S. van Ruiten, P. H. L. Krijger, H. Teunissen, R. H. Medema, B. van Steensel, T. R. Brummelkamp, E. de Wit, B. D. Rowland, The Cohesin Release Factor WAPL Restricts Chromatin Loop Extension. Cell 169, 693–707.e14 (2017).

60. P. Mach, P. I. Kos, Y. Zhan, J. Cramard, S. Gaudin, J. Tünnermann, E. Marchi, J. Eglinger, J. Zuin, M. Kryzhanovska, S. Smallwood, L. Gelman, G. Roth, E. P. Nora, G. Tiana, L. Giorgetti, Cohesin and CTCF control the dynamics of chromosome folding. Nat. Genet. 54, 1907–1918 (2022).

61. T.-H. S. Hsieh, A. Weiner, B. Lajoie, J. Dekker, N. Friedman, O. J. Rando, Mapping Nucleosome Resolution Chromosome Folding in Yeast by Micro-C. Cell 162, 108–119 (2015).

62. E. Oberbeckmann, K. Quililan, P. Cramer, A. M. Oudelaar, In vitro reconstitution of chromatin domains shows a role for nucleosome positioning in 3D genome organization. Nat. Genet. 56, 483–492 (2024).

63. E. G. Swanson, Y. Mao, B. J. Mallory, M. R. Vollger, S. C. Bohaczuk, C. B. Oliveira, D. B. Lyon, J. Ranchalis, N. L. Parmalee, B. A. Cohen, J. T. Bennett, A. B. Stergachis, Mapping single-cell diploid chromatin fiber architectures using DAF-seq. Nat. Biotechnol., 1–12 (2025).

64. R. He, W. Dong, Z. Wang, C. Xie, L. Gao, W. Ma, K. Shen, D. Li, Y. Pang, F. Jian, J. Zhang, Y. Yuan, X. Wang, Z. Zhang, Y. Zheng, S. Liu, C. Luo, X. Chai, J. Ren, Z. Zhu, X. S. Xie, Genome-wide single-cell and single-molecule footprinting of transcription factors with deaminase. Proc. Natl. Acad. Sci. 121, e2423270121 (2024).

65. B. V. Eckhardt, H. J. Richter, I. Rondeel, K. Renduchintala, F. Mattiroli, V. Ramani, The eukaryotic replisome intrinsically generates asymmetric daughter chromatin fibers. bioRxiv [Preprint] (2025). 10.1101/2025.09.18.677126.

66. A. G. Larson, D. Elnatan, M. M. Keenen, M. J. Trnka, J. B. Johnston, A. L. Burlingame, D. A. Agard, S. Redding, G. J. Narlikar, Liquid droplet formation by HP1α suggests a role for phase separation in heterochromatin. Nature 547, 236–240 (2017).

67. S. A. Quinodoz, N. Ollikainen, B. Tabak, A. Palla, J. M. Schmidt, E. Detmar, M. M. Lai, A. A. Shishkin, P. Bhat, Y. Takei, V. Trinh, E. Aznauryan, P. Russell, C. Cheng, M. Jovanovic, A. Chow, L. Cai, P. McDonel, M. Garber, M. Guttman, Higher-Order Inter-chromosomal Hubs Shape 3D Genome Organization in the Nucleus. Cell 174, 744–757.e24 (2018).

68. A. J. Plys, C. P. Davis, J. Kim, G. Rizki, M. M. Keenen, S. K. Marr, R. E. Kingston, Phase separation of Polycomb-repressive complex 1 is governed by a charged disordered region of CBX2. Genes Dev. 33, 799–813 (2019).

69. N. G. Aboreden, J. C. Lam, V. Y. Goel, S. Wang, X. Wang, S. C. Midla, A. Quijano, C. A. Keller, B. M. Giardine, R. C. Hardison, H. Zhang, A. S. Hansen, G. A. Blobel, LDB1 establishes multi-enhancer networks to regulate gene expression. Mol. Cell 85, 376–393.e9 (2025).

70. A. R. Strom, C. P. Brangwynne, The liquid nucleome – phase transitions in the nucleus at a glance. J. Cell Sci. 132, jcs235093 (2019).

71. B. J. E. Martin, E. F. Ablondi, C. Goglia, C. A. Mimoso, P. R. Espinel-Cabrera, K. Adelman, Global identification of SWI/SNF targets reveals compensation by EP400. Cell 186, 5290–5307.e26 (2023).

72. D. Barisic, M. B. Stadler, M. Iurlaro, D. Schübeler, Mammalian ISWI and SWI/SNF selectively mediate binding of distinct transcription factors. Nature 569, 136–140 (2019).

73. A. Jain, R. D. Vale, RNA phase transitions in repeat expansion disorders. Nature 546, 243–247 (2017).

74. G. K. Datar, E. Khabusheva, A. Anand, J. Beale, M. Sadek, C.-W. Chen, E. Potolitsyna, N. Alcantara-Contessoto, G. Liu, J. De La Fuente, C. Dollinger, A. Guzman, A. Martell, K. Wohlan, A. Maiti, N. J. Short, S. S. Yi, V. Andresen, B. T. Gjertsen, B. Falini, R. E. Rau, L. Brunetti, N. Sahni, M. A. Goodell, J. A. Riback, Disparate leukemia mutations converge on nuclear phase-separated condensates. Cell 188, 7118–7136.e21 (2025).

75. A. Bird, Cohesin as an essential disruptor of chromosome organization. Mol. Cell 85, 1054–1057 (2025).

76. J. Pulupa, N. G. McArthur, O. Stathi, M. Wang, M. Zazhytska, I. D. Pirozzolo, A. Nayar, L. Shapiro, S. Lomvardas, Solid phase transitions as a solution to the genome folding paradox. Nature 643, 820–829 (2025).

77. L. Kiefer, S. Gaudin, S. M. Rajkumar, G. I. F. Servito, J. Langen, M. H. Mui, S. Nawsheen, D. Canzio, Tuning cohesin trajectories enables differential readout of the Pcdhα cluster across neurons. Science 385, eadm9802 (2024).

78. L. Kiefer, A. Chiosso, J. Langen, A. Buckley, S. Gaudin, S. M. Rajkumar, G. I. F. Servito, E. S. Cha, A. Vijay, A. Yeung, A. Horta, M. H. Mui, D. Canzio, WAPL functions as a rheostat of Protocadherin isoform diversity that controls neural wiring. Science 380, eadf8440 (2023).

79. N. Q. Liu, M. Maresca, T. van den Brand, L. Braccioli, M. M. G. A. Schijns, H. Teunissen, B. G. Bruneau, E. P. Nora, E. de Wit, WAPL maintains a cohesin loading cycle to preserve cell-type-specific distal gene regulation. Nat. Genet. 53, 100–109 (2021).

80. Open2C, N. Abdennur, G. Fudenberg, I. M. Flyamer, A. A. Galitsyna, A. Goloborodko, M. Imakaev, S. V. Venev, Pairtools: From sequencing data to chromosome contacts. PLOS Comput. Biol. 20, e1012164 (2024).

81. N. Abdennur, L. A. Mirny, Cooler: scalable storage for Hi-C data and other genomically labeled arrays. Bioinformatics 36, 311–316 (2020).

82. Open2C, N. Abdennur, S. Abraham, G. Fudenberg, I. M. Flyamer, A. A. Galitsyna, A. Goloborodko, M. Imakaev, B. A. Oksuz, S. V. Venev, Y. Xiao, Cooltools: Enabling high-resolution Hi-C analysis in Python. PLOS Comput. Biol. 20, e1012067 (2024).

83. S. Tamburri, E. Lavarone, D. Fernández-Pérez, E. Conway, M. Zanotti, D. Manganaro, D. Pasini, Histone H2AK119 Mono-Ubiquitination Is Essential for Polycomb-Mediated Transcriptional Repression. Mol. Cell 77, 840–856.e5 (2020).

84. G. Fiorin, M. L. Klein, J. Hénin, Using collective variables to drive molecular dynamics simulations. Mol. Phys. 111, 3345–3362 (2013).

85. A. Grossfield, WHAM: the weighted histogram analysis method; http://membrane.urmc.rochester.edu/wordpress/?page_id=126.

86. A. J. C. Ladd, L. V. Woodcock, Triple-point coexistence properties of the lennard-jones system. Chem. Phys. Lett. 51, 155–159 (1977).

